# Sp Transcription Factors Establish the Signaling Environment in the Neuromesodermal Progenitor Niche During Axial Elongation

**DOI:** 10.1101/2025.06.03.657492

**Authors:** Ravindra B. Chalamalasetty, Haley Tran, Ryan Kelly, Samuel Kuo, Mark W. Kennedy, Moonsup Lee, Sara Thomas, Nikolaos Mandalos, Vishal Koparde, Francisco Pereira Lobo, Terry P. Yamaguchi

## Abstract

Neuromesodermal competent progenitors (NMPs) reside in the caudal epiblast near the node and primitive streak and give rise to spinal cord and somitic mesoderm during trunk and tail elongation. Their self-renewal depends on an autoregulatory loop involving Wnt3a and Fgf signaling, and the Tbxt and Cdx transcription factors, but the mechanisms underlying loop formation and the establishment of the niche are poorly understood. Here, we identify the zinc-finger transcription factors *Sp5* and *Sp8* (*Sp5/8*) as essential regulators of NMP maintenance. *Sp5/8* expression is controlled by Wnt, Fgf, Bmp and retinoic acid signaling, and they cooperate with Tbxt, Tcf7 and Cdx2 to sustain a robust autoregulatory network that promotes high Wnt/Fgf and low retinoic acid activity in the niche. These factors bind a novel enhancer essential for *Wnt3a* expression and feedback-loop integrity. Mechanistically, Sp5/8 regulates Tcf7 and Tle chromatin occupancy at Wnt-responsive enhancers suggesting their function as secondary Wnt transcriptional effectors. Our findings define a transcriptional module centered on Sp5/8 that stabilizes niche signaling and transcriptional circuitry essential for NMP fate decisions and trunk development.

## Introduction

The formation of the mammalian trunk and tail depends on a population of self-renewing axial progenitors located at the posterior end of the embryo, which continuously supply new cells for body axis elongation (Aires et al., 2018; Schnirman et al., 2023; Steventon and Martinez Arias, 2017; Wilson et al., 2009). Known as neuromesodermal competent progenitors (NMPs or NMCs; see Binagui-Casas et al., 2021 for discussion), they reside in the caudal lateral epiblast (CLE) near the primitive streak (PS) at the node-streak border (NSB), and contribute to both neural (spinal cord) and mesodermal (somite) lineages (Binagui-Casas et al., 2021; Cambray and Wilson, 2002, 2007; Garriock et al., 2015; Gouti et al., 2014; Henrique et al., 2015; Tzouanacou et al., 2009; Wymeersch et al., 2016). Their fate is determined by the relative expression of the transcription factors (TFs) Tbxt (Brachyury), which promotes mesoderm, and Sox2, which promotes neural identity (Garriock et al., 2015; Koch et al., 2017; Martin and Kimelman, 2012; Tsakiridis et al., 2014; Wymeersch et al., 2016). Co-expression of both factors maintains NMP bipotency (Wymeersch et al., 2016).

The canonical Wnt/β-catenin signaling pathway plays a central role in regulating *Tbxt* levels and, consequently, NMP fate. In the absence of Wnt ligands, β-catenin is degraded by the destruction complex, and repressive Tcf/Lef TFs such as Tcf7l1(Tcf3) inhibit Wnt target genes through interactions with Groucho/Tle co-repressors (Bou-Rouphael and Durand, 2021; Ramakrishnan and Cadigan, 2017). Wnt ligand binding to cell surface receptors of the Frizzled and LRP families stabilizes β-catenin, allowing its nuclear accumulation and activation of target genes such as *Tbxt* (Arnold et al., 2000; Galceran et al., 2001; Yamaguchi et al., 1999), via interaction with activating Tcfs such as Tcf7 and Lef1 at Wnt response elements (WREs).

Genetic studies demonstrate that coordinated Wnt, Fgf and Retinoic Acid (RA) signaling regulates *Tbxt* and *Sox2* expression. Loss of function (LOF) mutations in Wnt/β-catenin pathway components including *Wnt3a, Wnt8a, Ctnnb1* (β-catenin)*, Tcf7*, *Lef1,* or the target genes *Cdx1* and *2*, reduce *Tbxt* and increase *Sox2* levels, resulting in the loss of NMPs and the formation of ectopic neural tissue (Amin et al., 2016; Anand et al., 2023; Boulet and Capecchi, 2012; Cunningham et al., 2015; Dunty et al., 2008; Greco et al., 1996; Naiche et al., 2011; Takada et al., 1994; van de Ven et al., 2011; Wymeersch et al., 2016; Yamaguchi et al., 1999; Yoshikawa et al., 1997; Zhu and Lohnes, 2022). Similar phenotypes arise in *Fgf4/8* mutants (Boulet and Capecchi, 2012; Naiche et al., 2011) and in embryos lacking *Cyp26a1,* a RA-degrading enzyme expressed in NMPs under the control of Tbxt (Guibentif et al., 2021; Martin and Kimelman, 2010). *Cyp26a1* mutants display increased RA and *Sox2*, reduced *Wnt3a* and *Tbxt,* and impaired axial elongation (Abu-Abed et al., 2001; Abu-Abed et al., 2003; Sakai et al., 2001). Thus, a balance of Wnt, Fgf, and RA signals regulate NMP maintenance and differentiation.

Positive feedback loops further stabilize this signaling environment in the NMP niche. Wnt3a activates *Tbxt* via Tcf7 and Lef1, and in turn, Tbxt enhances *Wnt3a* expression, forming an autoregulatory loop critical for progenitor maintenance (Arnold et al., 2000; Galceran et al., 2001; Koch et al., 2017; Martin and Kimelman, 2012; Yamaguchi et al., 1999). Wnt-induced Cdx factors, especially Cdx2, reinforce this network by co-regulating Wnt and Fgf target genes (Amin et al., 2016; Young et al., 2009; Zhu and Lohnes, 2022), however the molecular basis of Tbxt**-**Cdx2**-**mediated regulation of *Wnt3a* remains poorly understood.

Adding further complexity to the Wnt/β-catenin transcriptional network, two zinc-finger TFs from the Specificity protein (Sp) family, Sp5 and Sp8, when mutated display axial truncations phenotypically similar to *Wnt3a* and *Tcf7/Lef1* mutants (Dunty et al., 2014; Galceran et al., 1999; Kennedy et al., 2016) suggesting functional interactions with canonical Wnt effectors. Both factors physically interact with Tcf7 and Lef1 (Kennedy et al., 2016), though their mechanistic roles are unclear. Here, we identify Sp5/8 as key transcriptional regulators in the NMP niche. We show that axial truncation in Sp5/8 mutants results from the loss of trunk NMPs, and that Sp5/8 directly regulate genes in the Wnt, Fgf, and RA pathways. Notably, we uncover a novel enhancer downstream of *Wnt3a* that is co-bound by Sp5/8, Tcf7, Tbxt, and Cdx2, and is required for *Wnt3a* expression. We further show that Sp5/8 stabilize Tcf7, while reducing Tle, occupancy at this enhancer, suggesting that Sp’s regulate the distribution of Tcf activator and repressive complexes. Together, our results define an Sp5/8-Tcf7-Tbxt-Cdx2 transcriptional module that sustains high Wnt and Fgf signaling and low RA activity in the NMP niche, thereby maintaining axial progenitor self-renewal and trunk and tail development.

## Results

### *Sp5* and *Sp8* are coexpressed in axial progenitors and are required for trunk development

*Sp8^Δ/Δ^* mutants die at birth with exencephaly, spina bifida and truncations of the tail and limbs, while *Sp5^LacZ/LacZ^* animals do not display a phenotype (Bell et al., 2003; Dunty et al., 2014; Harrison et al., 2000). Double null *Sp5^lacZ/lacZ^;Sp8^Δ/Δ^* mutants die earlier (∼E14) and display exencephaly, ectopic neural tubes caudally and more severe, i.e. more anterior, axial truncations, than *Sp8^Δ/Δ^* single mutants. Double mutants generate 12 somites and do not maintain *Tbxt* expression suggesting that *Sp5* and *Sp8* are required for trunk NMPs (Dunty et al., 2014). To assess a role for *Sp5* and *Sp8* specifically in NMPs, we used the *T-Cre* transgene, which targets axial trunk progenitors in the CLE and the PS but not the node (Garriock et al., 2015; Perantoni et al., 2005), to conditionally delete *Sp8^fl/fl^* in *Sp5^LacZ/LacZ^* embryos. *T-Cre*; *Sp5^LacZ/LacZ^*; *Sp8^fl/fl^* mutants (hereafter referred to as *T-Cre; Sp5/8 dko*) survived to birth, thereby circumventing the fetal lethality observed in *Sp5/8* double null mutants (Dunty et al., 2014). Analysis of *T-Cre; Sp5/8 dko* skeletons revealed that a few abnormal rostral ribs formed, but all skeletal elements caudal to mid-thoracic level T6 were absent (Figure 1A). The positioning of this axial truncation suggests that *Sp5*/*8* are redundantly required in trunk NMPs.

**Figure 1.**
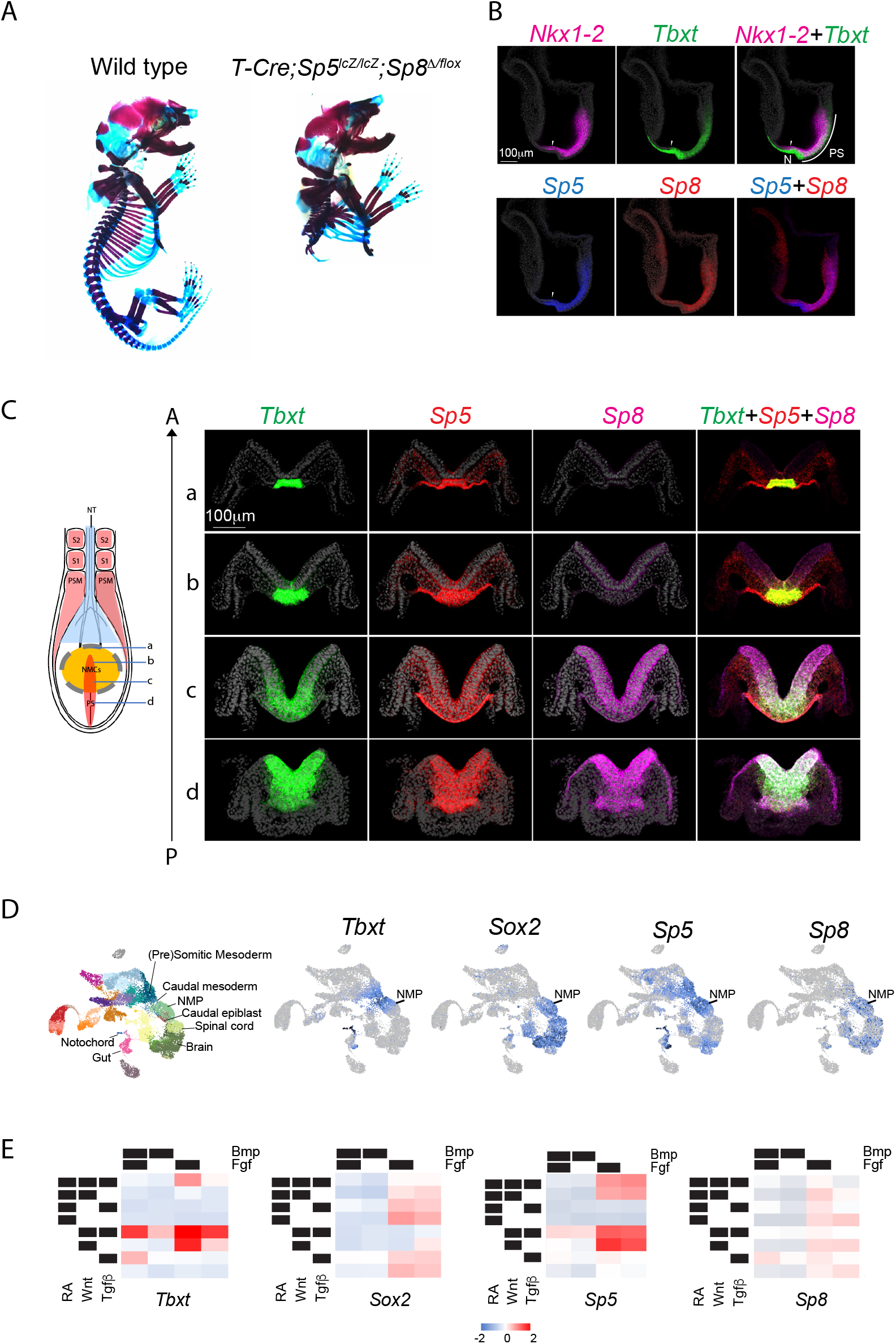
*Sp5* and *Sp8* are coexpressed in axial progenitors and are required for posterior development. A. Alcian blue (cartilage) and Alizarin red (bone) staining of E18.5 skeletons. B. Fluorescent in situ hybridization analysis of *Nkx1-2, Tbxt, Sp5* and *Sp8* expression in E7.75 wildtype embryos. Midline sagittal sections reveal that *Sp5* and *Tbxt* share an anterior boundary in the caudal epiblast (arrowhead). Abbreviations: N, node; PS, primitive streak. C. Whole mount fluorescent in situ hybridization analysis and optical cross sections of E8.5 wild type embryos for *Tbxt* (green), *Sp5* (Red), *Sp8* (purple) and Dapi (grey). The schematic on the left illustrates the axial levels at which optical sections were taken through the node (a), anterior PS/node streak border (b), mid primitive streak (C) and posterior PS (d). D. UMAP plots of single-cell RNAseq analysis of E8.25 mouse embryos showing *Tbxt, Sox2, Sp5,* and *Sp8* transcripts (Pijuan-Sala et al., 2019). E. Heatmaps depicting mean scaled gene expression of *Tbxt, Sox2, Sp5,* and *Sp8* in differentiating ESCs exposed to combinatorial Bmp, Fgf, RA, Wnt and Tgfβ signals (Yeo et al., 2020).

Fluorescent in situ hybridization chain reaction (HCR) (Choi et al., 2018) confirmed that *Sp5* and *Sp8* were coexpressed in trunk NMPs. At E7.75, both genes were expressed in the PS and caudal epiblast, overlapping extensively with the NMP markers *Tbxt* and *Nkx1-2* (Figure 1B). *Sp5’s* anterior expression boundary closely paralleled *Tbxt* in the epiblast near the NSB (arrowhead, Figure 1B), while *Sp8* expression overlapped with the pre-neural *Nkx1-2*-positive region (Rodrigo Albors et al., 2018) and extended anteriorly into the neural plate (Figure 1B) (Dunty et al., 2014). Optical sections revealed that *Sp5* and *Sp8* were coexpressed with *Tbxt* in the medial PS and hindgut at E8.5 but extended further laterally into the epiblast beyond the *Tbxt* domain (Figure 1C). Single cell RNA-seq analysis of E8.25 embryos (Pijuan-Sala et al., 2019) corroborated the HCR findings. *Sp5* expression mirrored *Tbxt* in NMPs, caudal and presomitic mesoderm, notochord and gut, while *Sp8* closely aligned with *Sox2* in NMPs and neural progenitors of the spinal cord and brain (Figure 1D). *Sp5* and *Sp8* were coexpressed in NMPs, supporting a dual role in trunk NMP maintenance.

To identify upstream regulators of *Sp5/8*, we explored signaling pathways known to influence NMP specification in vitro. *Sp5* and *Tbxt* were robustly activated in differentiating ESCs (Yeo et al., 2020) by combinatorial mesoderm-promoting (Wnt, Fgf) signals (Koch et al., 2017; Sudheer et al., 2016), but inhibited by RA, a neural inducer, or Bmp which promotes lateral mesoderm fates from NMP-derived mesoderm (Row et al., 2018) (Figure 1E). In contrast, *Sp8* was activated by neural-promoting (RA, Fgf) and mesoderm-promoting (Wnt, Fgf) signals but also repressed by Bmp (Figure 1E). Thus, *Sp5* and *Sp8* expression are distinctly, yet coordinately, regulated by signals that control NMP specification and differentiation.

### Transcriptional profiling suggests roles for Sp5/8 in the regulatory circuitry of NMPs

To identify genes co-regulated by *Sp5* and *Sp8*, bulk RNA-seq was performed on the posterior region of *Sp5/8* double-null embryos at E8.5 (4-6 somite stage (ss)), prior to overt morphological changes (Figure 2-figure supplement A, B). Over-Representation Analysis (ORA) of 564 downregulated genes revealed that these genes are associated with canonical Wnt signaling, gastrulation, mesoderm formation, somite development, and axis elongation (Figure 2A, Figure 2-figure supplement C, D, Supplementary file 1). These include TFs known to control NMP specification *(Tbxt, Cdx1, 2, 4*), posterior identity *(Hoxa9, Hoxb9, Hoxb7* and others), and presomitic mesoderm (PSM) differentiation and segmentation *(Lef1, Hes7, Tbx6, and Msgn1*) (Figure 2B). Additionally, key signaling pathways required for the development of axial progenitors – Wnt/β-catenin *(Wnt3a, Wnt5b, Axin2, Lef1, Cdx1,2,4, Tbxt, Msgn1)*, Fgf (*Fgf4, Fgf8, Fgf17, Dusp4, Etv4)*, Nodal (*Nodal, Lefty1, Lefty2*), Notch (*Notch1, Dll1, Dll3, Hes7*) and Bmp *(Gdf11*) – were disrupted in mutants (Aires et al., 2019; Dunty et al., 2014; Gouti et al., 2017; Jurberg et al., 2013; Koch et al., 2017; Wymeersch et al., 2019). Many affected genes, notably *Wnt3a, Tbxt*, *Cdx1* and *Fgf4*, as well as *Sp5* and *Sp8,* were bound by Sp5/8 in ChIP-seq assays performed in differentiating EBs (Figure 2C, 3B, Supplementary file 1), suggesting that Sp5 and Sp8 are autoregulatory, and participate directly in the Wnt3a-Tbxt-Cdx positive feedback loop.

**Figure 2.**
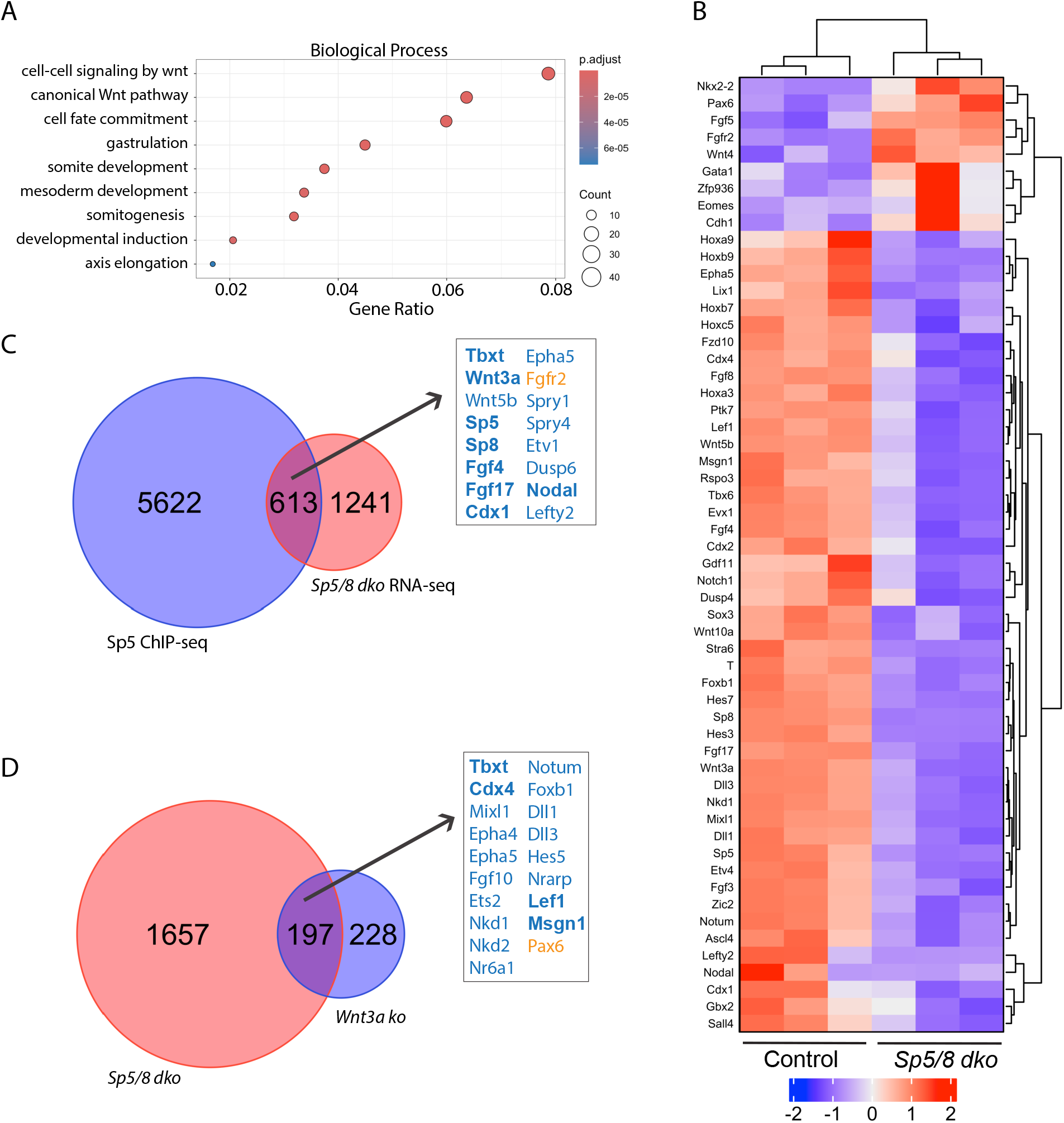
Sp5/8 regulates the Wnt3a/β-catenin pathway. A. Over Representation Analysis (ORA) of 564 downregulated genes in E8.5 *Sp5/8 dko* mutants. Color scale reflects adjusted P-value, circle size represents gene number. B. Heat map of log2 z-scores of normalized counts showing differentially expressed genes (P≤0.05; FC ≥0.5). C. Venn diagram illustrating the intersection between Sp5 ChIP-seq and *Sp5/8 dko* RNA-seq analyses. 613 differentially expressed genes, including *Wnt3a* and *Tbxt,* are direct targets of Sp5. Downregulated genes are depicted in blue and upregulated genes in orange. D. Venn diagram showing 197 differentially expressed genes in common between the *Sp5/8 dko* and *Wnt3a ko* transcriptomes. Downregulated genes (blue), upregulated genes (orange).

**Figure 3.**
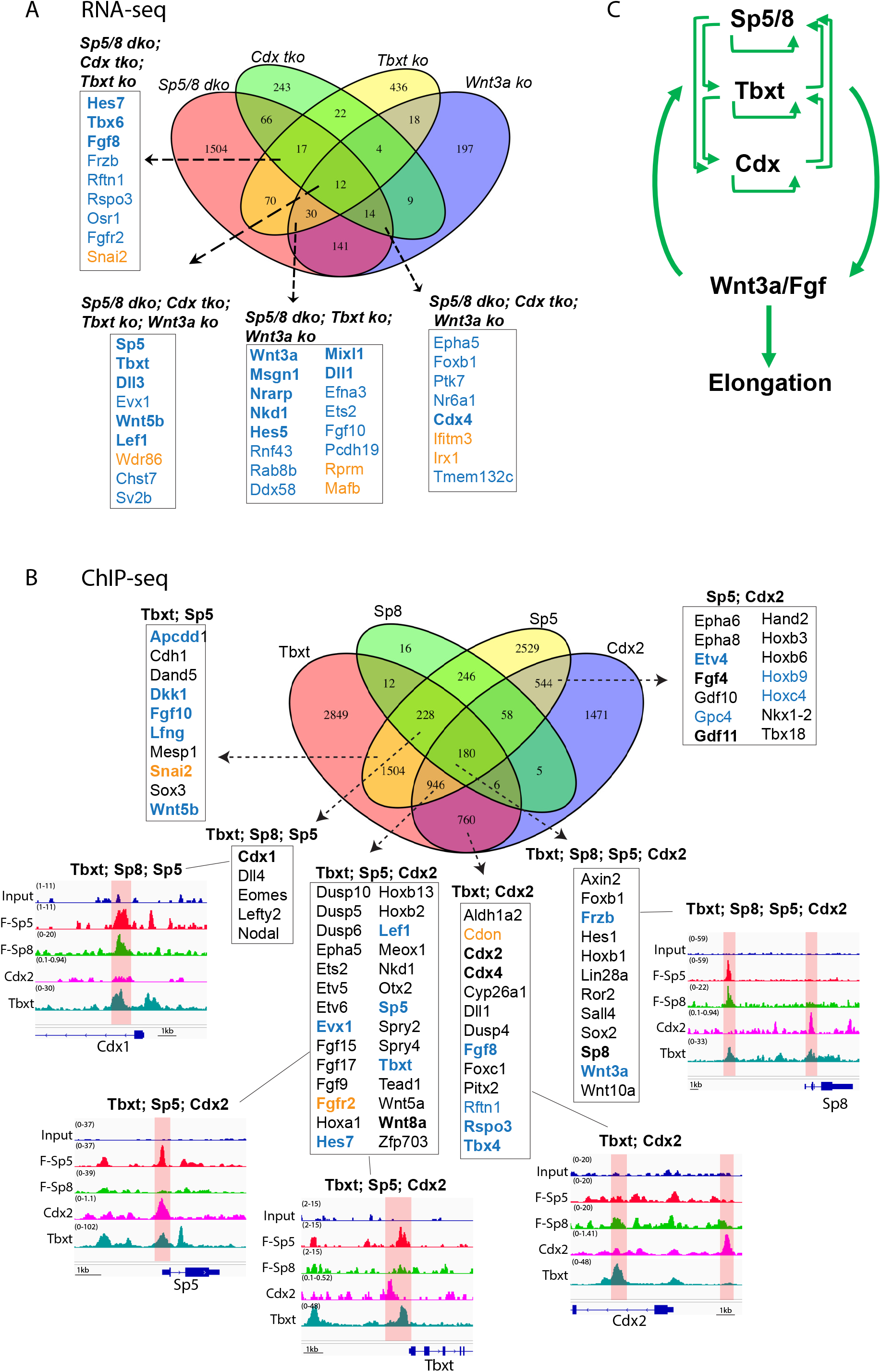
Sp5/8 are integral components of the core regulatory circuitry of axial progenitors. A. Four-way Venn diagram illustrating differentially expressed genes common to *Sp5/8 dko, Cdx tko* (Amin et al., 2016)*, Tbxt ko* (Koch et al., 2017), and *Wnt3a ko* transcriptomes. Key regulators are highlighted in bold, downregulated genes (blue), upregulated genes (orange). B. Four-way Venn diagram illustrating direct target genes bound combinatorially by Tbxt, Flag-Sp5, Flag-Sp8, and/or Cdx2. ChIP-seq was performed in D3 EBs for Flag-Sp5 and Flag-Sp8 (Kennedy et al., 2016), NMP populations differentiated in vitro from CHIR-treated ESCs for Tbxt or, in the case of Cdx2, EpiSCs (Amin et al., 2016). ChIP-seq tracks for a few key NMP genes are shown. Genes bound by the indicated TF and differentially expressed in the respective transcriptomes are color-coded blue (downregulated) or orange (upregulated). C. Sp5/8, Tbxt and Cdx TFs form self-sustaining autoregulatory loops that collectively maintain a Wnt3a/Fgf positive feedback loop to drive NMP self-renewal and axial elongation.

We reasoned that if Sp5 and Sp8 function in the Wnt3a-Tbxt-Cdx feedback loop then comparative RNA-seq analyses of E8.5 *Sp5/8*, *Wnt3a*, *Tbxt* (Koch et al., 2017) and *Cdx1/2/4* (Amin et al., 2016; Zhu and Lohnes, 2022) mutants should reveal common differentially expressed genes (DEGs). A comparison of *Sp5/8* and *Wnt3a* mutant transcriptomes identified a set of 197 genes (68% concordant), including the established Wnt targets *Tbxt, Cdx4, Lef1* and *Msgn1* (Figure 2D; 3A, Figure 2-figure supplement E, Supplementary file 2) (Chalamalasetty et al., 2014; Galceran et al., 2001; Hovanes et al., 2001; Yamaguchi et al., 1999). Notably, *Wnt3a* and *Sp5*/*8* displayed codependence as *Wnt3a* expression was reduced in *Sp5/8 dko* embryos (−2.93 log2FC), while *Sp5* (−1.9 log2FC) and *Sp8* (−0.92 log2FC) levels were reduced in *Wnt3a^-/-^*mutants. A similar codependency was observed between *Sp5/8* and the *Cdx* genes (Figure 3-figure supplement 1A, B), and *Sp5/8* and *Tbxt* (Figure 3-figure supplement 1C, D), when the *Sp5/8 dko* was compared to *Cdx tko* and *Tbxt ko* transcriptional profiles (Supplementary file 3). Further comparisons across all four mutant transcriptomes (*Sp5/8, Wnt3a, Tbxt* and *Cdx1/2/4)* identified a core set of 12 shared DEGs that included *Sp5, Wnt3a, Tbxt,* and *Lef1* (Figure 3A, Figure 3-figure supplement 1, Supplementary file 3). Examination of Sp5, Sp8, Tbxt and Cdx2 (Amin et al., 2016) ChIP-seq datasets, derived from in vitro-differentiated NMP populations, confirmed that cis-regulatory elements at *Sp5, Sp8, Tbxt, Cdx1, 2,* and *4* were bound directly by Tbxt and either Cdx2, Sp5, or Sp8 (Figure 3B, Supplementary file 4). The high degree of interaction between these TFs suggests that Sp5 and Sp8 are integral components of the regulatory circuitry of axial progenitors (Figure 3C).

### NMP and PSM fates depend on Sp5/8 activity in vivo

To characterize the role of Sp5 and Sp8 in the Wnt3a-Tbxt feedback loop and in trunk axial progenitors in vivo, we first examined the expression of the NMP-defining TFs Tbxt and Sox2 in *Sp5/8 dko* embryos at the 4-7ss stage when trunk somites are emerging and when axial progenitors are being generated. Whole-mount immunofluorescence (IF) showed that Tbxt was markedly reduced or absent from the anterior PS, while Sox2 was elevated and posteriorly expanded in mutants (Figure 4b), compared to controls (Figure 4a). Mid-sagittal sections confirmed that Tbxt+; Sox2+ NMPs were largely found at the NSB (Figure 4d-f), and that these double positive NMPs were absent in *Sp5/8 dko* mutants (Figure 4g-i). The low Tbxt/high Sox2 levels are consistent with an NMP-to-neural fate conversion in the absence of Sp5/8. In contrast, Sp8 overexpression in axial progenitors dramatically expanded the Tbxt expression domain, promoting overgrowth of the entire posterior region (Figure 4c; Figure 4-figure supplement 1A). These results show that Tbxt and Sox2 levels are controlled by Sp5 and Sp8.

**Figure 4.**
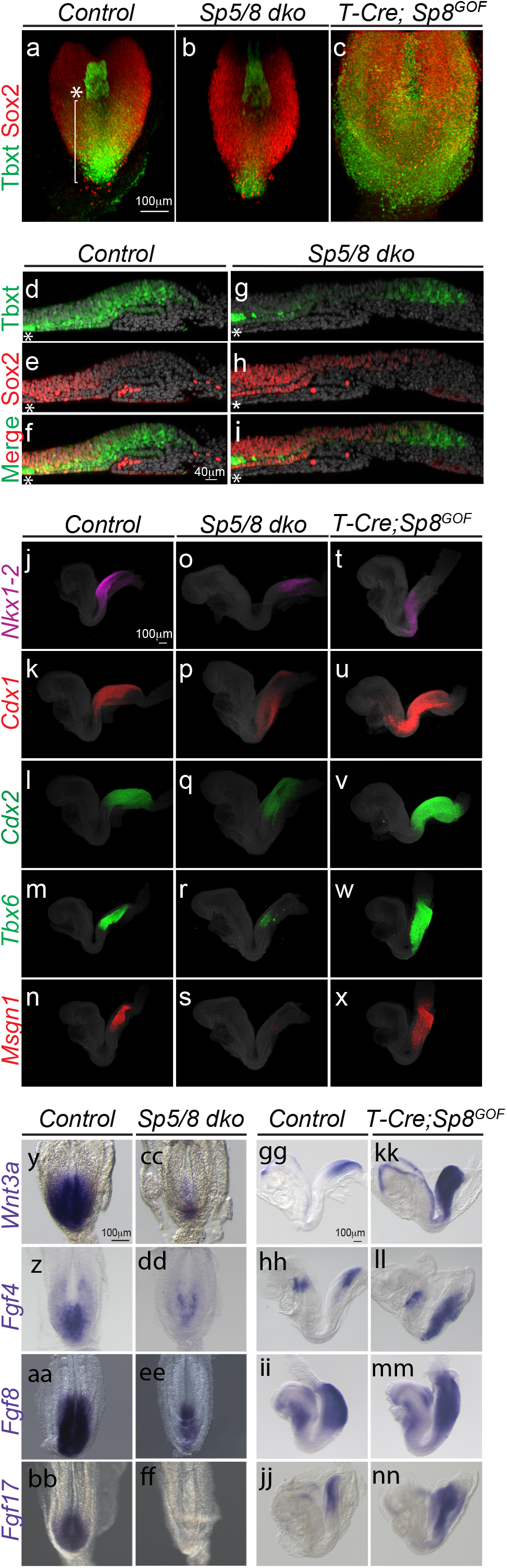
Sp5/8 regulate axial progenitors and the expression of niche factors. A-C. Projection images of whole mount immunofluorescence analysis showing Tbxt (green) and Sox2 (red) protein expression in the posterior region of E8.5 control (a), *Sp5/8 dko* (b) and *T-cre; Sp8^GOF^* (c) mutants. The bracket indicates the length of the primitive streak; the asterisk aligns with the caudal node. D-I. Midline sagittal sections taken through the primitive streak of flat mounted E8.5 control (d-f) and *Sp5/8 dko* embryos (g-i) processed for Tbxt (green), and Sox2 (red) IF (Dapi, grey). J-X. HCR analysis of *Nkx1-2* (magenta), *Cdx1* (red), *Cdx2* (green), *Tbx6* (green), and *Msgn1* (red) expression in E8.5 control (j-n), *Sp5/8 dko* (o-s) and *T-cre; Sp8^GOF^* (t-x) embryos. Y-NN. Chromogenic WISH of E8.5 control (y-bb) and *Sp5/8 dko* (cc-ff), E8.5 control (gg-jj) and *T-cre; Sp8^GOF^* (kk-nn) with DIG-labeled *Wnt3a (y, cc, gg, kk), Fgf4 (z, dd, hh, ll), Fgf8 (aa, ee, ii, m)* and *Fgf17 (bb, ff, jj, nn)* riboprobes. (y-ff) dorsal views, (gg-nn) lateral views.

HCR analysis of NMP (*Nkx1.2, Wnt8a)*, posterior identity (*Cdx1, Cdx2*), and PSM markers (*Tbx6, Msgn1*) (Amin et al., 2016; Cunningham et al., 2015; Garriock et al., 2020; Gouti et al., 2017; Rodrigo Albors et al., 2018; Zhu and Lohnes, 2022) revealed their reduction or absence in *Sp5/8* mutants, reinforcing the requirement for Sp5/8 in NMP and PSM maintenance (Figure 4j-s; Figure 4-figure supplement 1C). Interestingly, overexpression of Sp8 in axial progenitors resulted in reduced expression of the NMP markers *Nkx1.2* and *Wnt8a,* and a dramatic increase in the levels and spatial domains of the Wnt targets *Cdx1, Cdx2, Tbx6* and *Msgn1,* indicating that elevated Sp5/8 activity drives the Wnt pathway and posterior identity, and pushes NMP differentiation towards mesodermal fates (Figure 4-figure supplement 1B, D: Figure 4t-x). Together, the data suggest that Sp5/8 control NMP fate decisions in a dose-dependent manner; loss of Sp activity disrupts NMP maintenance, promoting neural fate, intermediate levels maintain NMPs, and high Sp5/8 levels favor mesoderm differentiation.

### Sp5/8 regulate the signaling environment of the NMP niche

RNA-seq analysis indicated disrupted expression of key genes in the Wnt, Fgf and RA signaling pathways in *Sp5/8* mutants, suggesting that Sp5/8 could control NMP fate decisions by regulating the niche (Figure 2, 3). WISH analysis confirmed that *Wnt8a* (Figure 4-figure supplement 1C)*, Wnt3a, Fgf4, Fgf8* and *Fgf17,* expressed in the CLE and PS of controls (Figure 4y-bb) (Boylan et al., 2020; Crossley and Martin, 1995; Cunningham et al., 2015; Yamaguchi and Rossant, 1995) were significantly reduced or absent in *Sp5/8* mutants (Figure 4cc-4ff). Additionally, expression of *Cyp26a1,* which degrades RA, was also downregulated, while *Aldh1a2* (an RA synthesis enzyme) levels were maintained, implying elevated levels of RA in mutant axial progenitors (Figure 4-figure supplement 2a-f). Collectively, these results show that loss of Sp activity disrupts the balance of morphogens - lowering Wnt and Fgf while increasing RA - thereby impairing NMP self-renewal.

Transient overexpression of Sp8 in axial progenitors led to elevated levels and expanded expression domains of *Wnt3a, Fgf4, Fgf8* and *Fgf17*, consistent with activation of the Wnt3a-Tbxt feedback loop (Figure 4gg-4nn). Fgf4 and 8 play an important role during axial elongation to maintain the undifferentiated state of presomitic progenitors (Boulet and Capecchi, 2012; Naiche et al., 2011). The increased and expanded expression of *Fgf4* and *8* predicts that the PSM progenitor pool was expanded at the expense of somitic differentiation. Increased expression of the PSM markers *Tbx6* and *Msgn1* (Figure 4w, x), coupled with the reduced expression domains of the somite markers *Uncx4.1* and *Aldh1a2* (Figure 4-figure supplement 1B, 4c, d), is consistent with a role for Sp8 in regulating the PSM progenitor pool through these Fgfs. These results are consistent with Sp5/8 functioning as regulators of niche signals critical for NMP fate decisions.

### An Sp5/8-Tcf-Cdx-Tbxt TF module regulates *Wnt3a* via a novel downstream enhancer

Given the crucial role that Wnt3a plays in NMP self-renewal, we focused our attention on understanding how Sp5/8 regulate *Wnt3a*. Examination of the ChIP-seq data for Sp5 and Sp8 binding sites led to the identification of peaks in a conserved, intergenic region 4.2 kb downstream of *Wnt3a* (Figure 5A; Figure 5-figure supplement 1A). This region bears the hallmarks of an enhancer as it is bound by H3K27ac, an epigenetic mark of active enhancers, in the PS and mesendoderm of E7.5 embryos when *Wnt3a* is first expressed in these tissues, but not in extraembryonic or earlier-staged tissues where it is not expressed (Xiang et al., 2020; Yang et al., 2019) (Figure 5B). Furthermore, scATAC-seq data from E8.5 mouse embryos showed that while the *Wnt3a* promoter was broadly accessible across all cell types, the putative enhancer element was accessible only in *Wnt3a-*expressing cells (Figure 5C) (Argelaguet et al., 2022; Pijuan-Sala et al., 2019). This included caudal epiblast, PS, NMP, caudal and somitic mesoderm, and notochord, but not neural, surface ectoderm, and gut which do not express *Wnt3a* at this stage. These findings suggest that this element regulates tissue-specific expression of *Wnt3a*. Hereafter, we refer to this element as the *Wnt3a-DE* (Downstream Enhancer).

**Figure 5.**
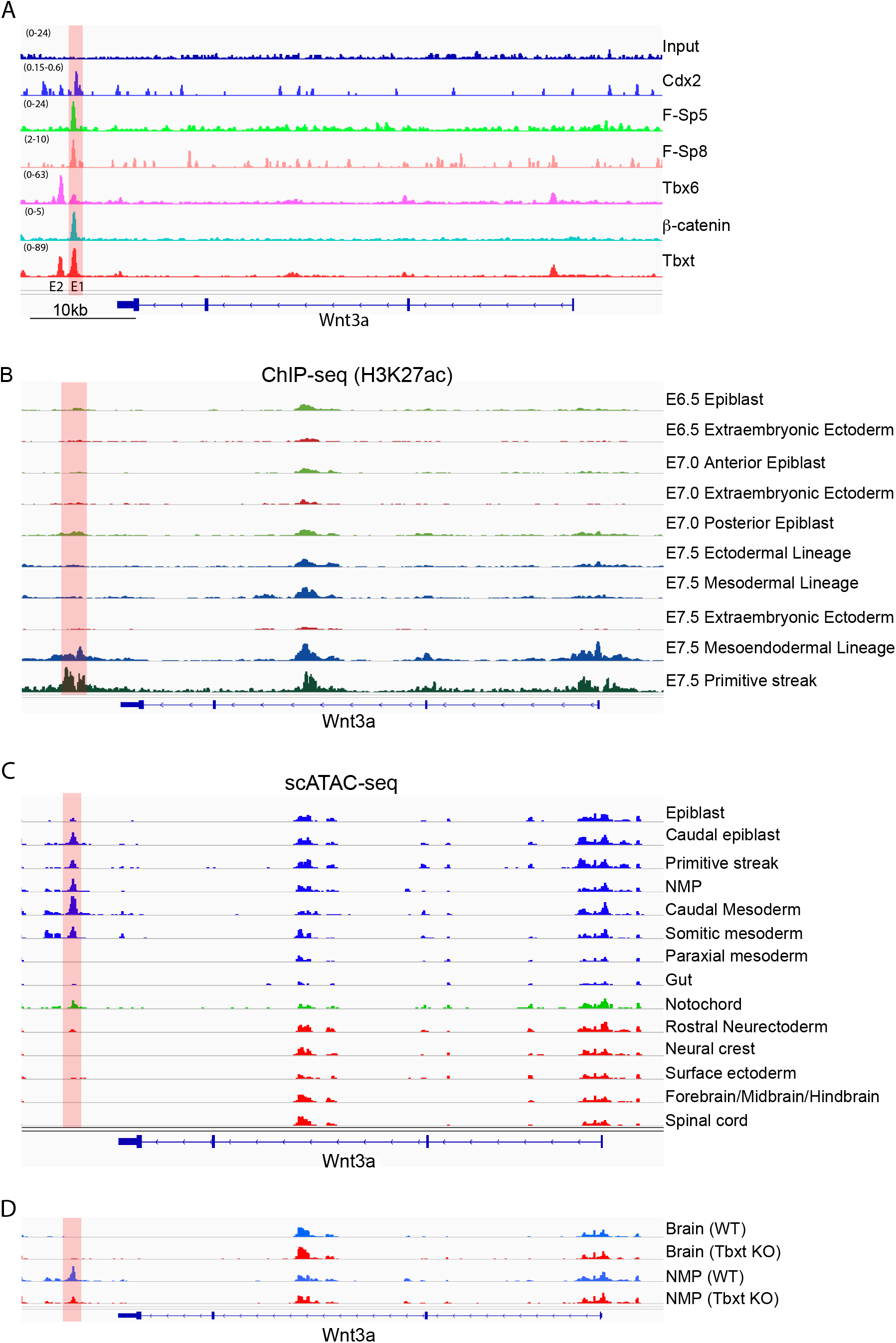
Identification of a novel *Wnt3a* downstream enhancer. A. Integrative Genomics Viewer (IGV) analysis of ChIP-seq peaks for Cdx2 (Amin et al., 2016), Flag-Sp5, Flag-Sp8, Tbx6 (Koch et al., 2017), β-catenin (Blassberg et al., 2022), Tbxt and Input negative control at *Wnt3a*. Two putative downstream enhancers are denoted as E1 (red bar, referred to as *Wnt3a^DE^* in text) and E2. B. ChIP-seq tracks derived from (Xiang et al., 2020) and (Yang et al., 2019) showing deposition of the active regulatory mark H3K27ac at both the promoter and putative enhancer region of *Wnt3a* in the mesendodermal and primitive streak lineages at E7.5. C. scATAC-seq data of E8.5 wild type mouse embryos showing accessibility across *Wnt3a* (Argelaguet et al., 2022). D. scATAC-seq data from wild type (WT) and *Tbxt* knockout (KO) mouse embryos at E8.5 showing reduced accessibility at *Wnt3a^DE^* in the absence of *Tbxt* (Argelaguet et al., 2022).

Further mining of ChIP-seq datasets showed that in addition to binding Sp5 and Sp8, the *Wnt3a-DE* is co-occupied by the NMP regulators β-catenin, Cdx2 and Tbxt but not Tbx6 (Figure 5A), and contains consensus binding motifs for Sp5, Tcf/Lef, Cdx and Tbx, suggesting direct regulation by these TFs (Figure 5-figure supplement 1B). To examine the functional consequences of Tbxt binding to the *Wnt3a-DE*, we analyzed scATAC-seq data performed on E8.5 *Tbxt^-/-^* embryos (Argelaguet et al., 2022). Loss of *Tbxt* resulted in reduced accessibility of the *Wnt3a-DE* in NMPs (Figure 5D) suggesting that Tbxt regulates *Wnt3a* expression in NMPs by controlling chromatin accessibility.

Functional characterization of the *Wnt3a-DE* (Enhancer E1) and E2 elements (Figure 5A) in luciferase reporter assays in differentiating ESCs confirmed that the Wnt3a-DE, but not the E2 element, could drive transcription in response to CHIR, a GSK3 inhibitor, and bFgf treatment (Figure 6A). Combining the E2 element with Wnt3a-DE reduced the activity of Wnt3a-DE by 10- fold suggesting that E2 possesses silencer activity, potentially mediated by Tbx6. Overexpression of Flag-Sp5 further enhanced transcriptional activity of the Wnt3a-DE element when stimulated by CHIR alone or together with bFgf (Figure 6B). Enhancer deletion experiments showed that the Sp and Cdx sites are necessary but not sufficient for enhancer activity (Figure 6C). Similarly, Tcf/Lef and Tbxt binding sites are necessary but insufficient for enhancer activity suggesting that all four TFs are required for robust Wnt3a-DE activity.

**Figure 6.**
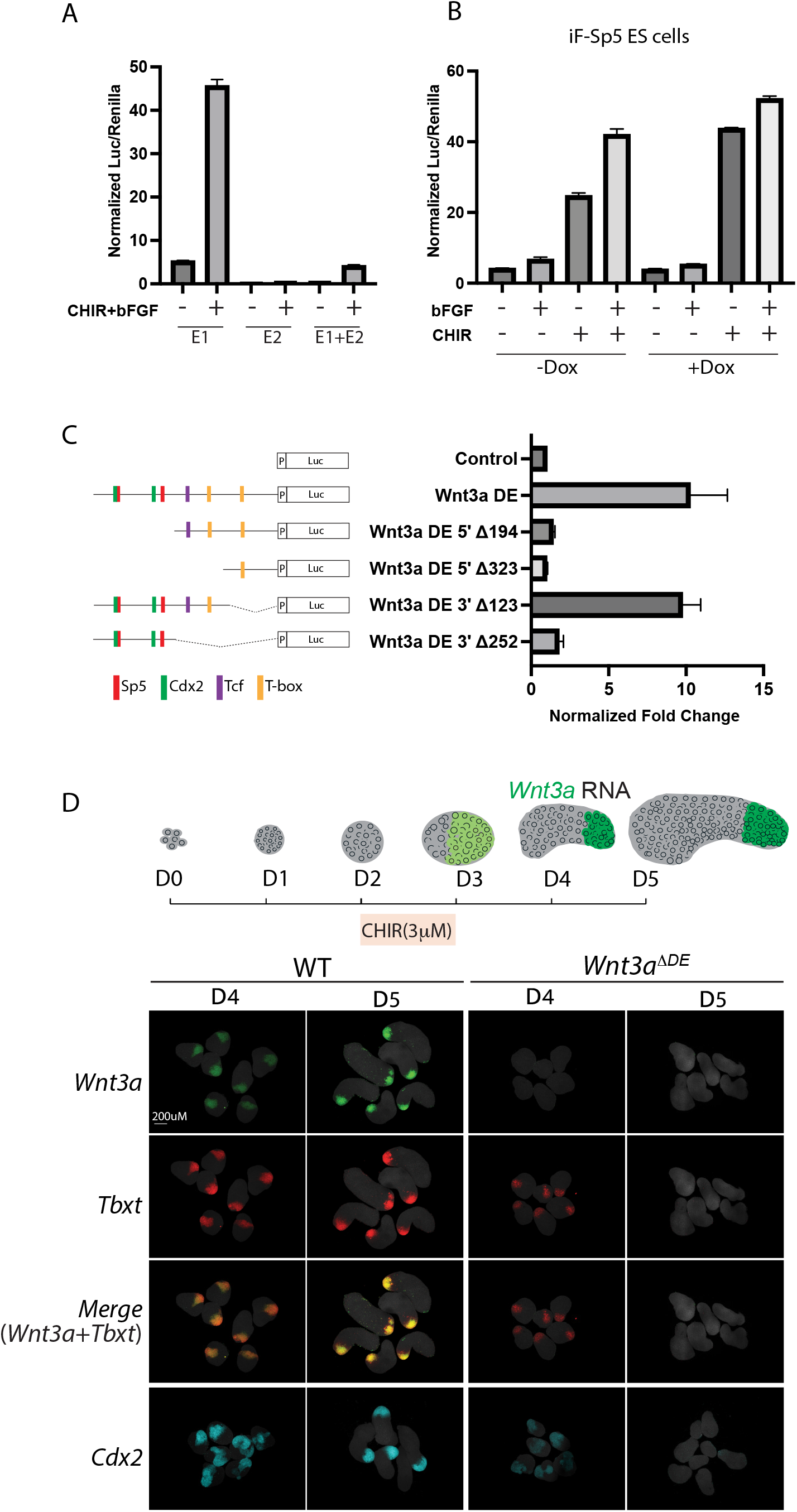
The *Wnt3a^DE^* is a WRE necessary for expression of *Wnt3a* and the *Wnt3a-Tbxt* autoregulatory loop. A. Luciferase reporter assays of *Wnt3a* enhancer constructs: 0.47kb E1 (*Wnt3a^DE^*), 0.8kb E2 and the 1.8kb E1+E2, in differentiating ESCs treated with CHIR and Fgf to generate NMPs. Data are depicted as Mean+/- SD (n=3). Y-axis, relative luciferase units. B. Sp5 overexpression enhances *Wnt3a^DE^*-luc activity in differentiating i3xFlag-Sp5 ES cells treated with bFgf (12ng/ml), CHIR (3µM), or bFgf+CHIR +/- Dox to induce 3xFlag-Sp5. C. Luciferase assays of *Wnt3a^DE^*-luc deletion constructs. Schematic depicts the relative locations of Sp5, Cdx2, Tcf/Lef, and T-box binding motifs in the deletion constructs. D. (Top) Schematic showing polarized *Wnt3a* expression (green) in elongating gastruloids. (Bottom) HCR analysis of *Wnt3a* (green), *Tbxt* (red), and *Cdx2* (cyan) expression in WT(R1) and *Wnt3a^ΔDE^* gastruloids one (D4) and two (D5) days after CHIR treatment, respectively. Dapi, grey.

To determine if Wnt3a-DE is necessary for *Wnt3a* expression during NMP formation and axial elongation, we deleted the element in ESCs using CRISPR/Cas9 gene editing (Figure 6-figure supplement 1A) and examined the consequences of this mutation on gastruloid development. Gastruloids are self-organizing three-dimensional ESC aggregates that undergo morphogenesis in vitro characteristic of embryo axial elongation (Beccari et al., 2018; van den Brink et al., 2020; Veenvliet et al., 2020). A brief treatment of epiblast-like aggregates with CHIR initiates PS differentiation programs that establish the Wnt3a-Tbxt autoregulatory loop that leads to posterior polarization and axial elongation. HCR WISH demonstrated that posterior-polarized domains of *Wnt3a, Tbxt* and *Cdx2* co-expression were apparent in control gastruloids by D4 and highly expressed by D5 due to positive feedback (Figure 6D). In contrast, gastruloids lacking the *Wnt3a- DE* (*Wnt3a^ΔDE^*) did not express *Wnt3a* and only expressed low levels of *Tbxt* and *Cdx2* on D4 that were not maintained on D5 (Figure 6D, right). Sox2 expression remained unchanged between control and *Wnt3a^ΔDE^* gastruloids (Figure 6-figure supplement 1B). Thus, the *Wnt3a-DE* is necessary for *Wnt3a* expression and to sustain the Sp5/8-Tcf-Cdx-Tbxt autoregulatory loop that maintains NMPs.

### Sp5 regulates Wnt target gene expression by controlling Tcf7 and Tle occupancy at WREs

We have previously shown protein-protein interactions between Sp5/8 and Tcf7/Lef1 (Kennedy et al., 2016), however the functional relevance of this interaction for Tcf7/Lef1 activity and Wnt target gene transcription remains unclear. To determine whether Sp5/8 regulate Tcf7 binding at WREs, we performed ChIP-seq for Tcf7 and Flag-Sp5 in CHIR-stimulated embryoid bodies (EBs) generated from WT or *Sp5/8 dko* ESCs (Figure 6-figure supplement 1C, D; Figure 6-figure supplement 1C, D source data 1). 28,154 Tcf7 peaks were identified (Figure 7-figure supplement 1A, Supplementary file 5). Interestingly, the average Tcf7 peak intensity value, which is a measure of read density at a specific genomic region, was reduced by half in *Sp5/8 dko* cells, suggesting that Sp TFs regulate Tcf7 binding. Examination of genomic regions that bound both Tcf7 and Flag-Sp5 identified 3471 sites, corresponding to 1168 genes, that were co-occupied by both TFs (Figure 7-figure supplement 1B). The average Tcf7 peak intensity value at these putative WREs was considerably higher (35) than that observed for genomic Tcf7 binding sites without a demonstrated Sp5 peak (17) (cf. Figure 7A and Figure 7-figure supplement 1A) suggesting that the presence of Sp5 enhanced the binding of Tcf7 to DNA. This is supported by the observation that Tcf7 binding strength at these co-occupied sites was reduced by ∼30% in *Sp5/8 dko* cells (Figure 7A). This reduction in Tcf7 binding was not due to changes in Tcf7 expression as Tcf7 protein and RNA levels did not depend on Sp5/8 levels (Figure 7-figure supplement 1C, D). We then asked whether these changes in Tcf7 binding activity correlated with changes in Wnt target gene transcription. Of the 1168 genes co-bound by Tcf7 and Sp5, 81 were downregulated in *Sp5/8 dko* embryos, including the well-characterized Wnt/Sp target genes *Cdx1,2, Msgn1, Tbxt* and *Tbx6* (Figure 7-figure supplement 1B). Together, our findings suggest that Sp5/8 modulate Wnt target gene transcription by enhancing Tcf7 occupancy at WREs.

**Figure 7.**
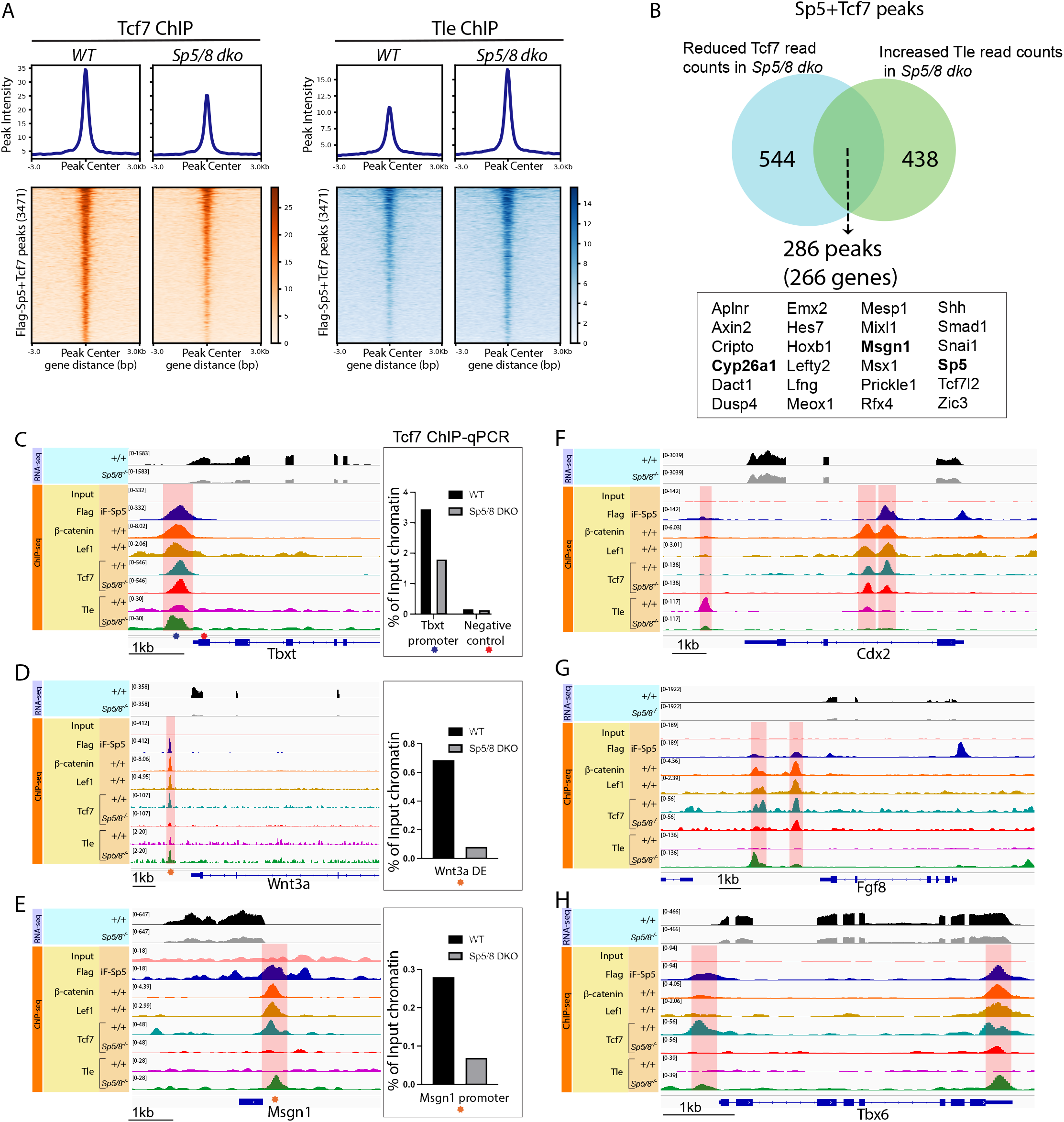
Sp5/8 regulate Tcf7 and Tle occupancy at WREs. A. Metaplots and heatmaps of Tcf7 and Tle binding profiles at sites of Sp5 and Tcf7 co-occupancy in WT and *Sp5/8 dko* NMPs. B. Identification of potential sites of Sp5/8-dependent Tcf exchange. 286 WREs defined by Sp5 and Tcf7 co-occupancy in wt cells display reduced Tcf7, and increased Tle, peak intensities in *Sp5/8 dko* cells. Only peaks with a log2 fold change > 0.58 and a P-value ≤ 0.05 were included in this analysis. C. IGV genome tracks of RNA-seq and ChIP-seq at select Wnt target genes in +/+ and *Sp5/8^-/-^* (*dko*) mutants. Top rows in each set depict RNAseq expression data obtained from E8.5 +/+ and *Sp5/8^-/-^* embryos. Bottom rows illustrate ChIP-seq data for Input, Flag-Sp5 (in iF-Sp5 cells), Tcf7 and Tle performed in +/+ and *Sp5/8^-/-^* EBs. β-catenin, Lef1 ChIP-seq data from (Blassberg et al., 2022). *Tbxt* (a)*, Wnt3a* (b)*, Msgn1* (c), *Cdx2* (d)*, Fgf8* (e) and *Tbx6* (f) loci are shown. ChIP-qPCR to validate Tcf7 occupancy is shown on the right for *Tbxt* (c)*, Wnt3a* (d), and *Msgn1* (e).

In mouse embryos and ESCs, *Tcf7l1*(Tcf3) functions in the epiblast as a dedicated repressor of Wnt target genes while Tcf7 and Lef1 activate these targets (Galceran et al., 1999; Hoffman et al., 2013; Wu et al., 2012; Yi et al., 2011). This is consistent with a “Tcf exchange” model in which repressive Tcf7l1-Tle complexes bind WREs to suppress Wnt target genes in the absence of Wnt signaling (i.e. the OFF state), and are exchanged for activating Tcf7/Lef1-β-catenin complexes to initiate target gene expression in the presence of Wnt (i.e. the ON state) (Guo et al., 2021; Ramakrishnan et al., 2018). Alternatively, Tcf7-Tle bound to Wnt target genes in the OFF state could switch from a repressor to an activator upon binding of β-catenin (van Tienen et al., 2017). To examine the genomic distribution of Tle co-repressors in the presence or absence of Sp5/8, a pan-Tle antibody was chosen for ChIP as it detected Tle1-4 regardless of their binding partners. We then asked if Tle1-4 could be detected at the 3471 Tcf7-Sp5 co-bound WREs. Small Tle peaks were identified at these WREs in WT cells (Figure 7A). Intriguingly, whereas Tcf7 peak intensity was reduced at WREs in *Sp5/8 dko* cells, Tle peak intensities nearly doubled (Figure 7A, Figure 7-figure supplement 1A). qPCR and Western blot experiments suggested that increased Tle1-4 binding was not due to elevated Tle1-4 expression since no significant changes in protein or RNA levels were detected in the presence or absence of Sp5/8 (Figure 7-figure supplement 1C, D; Figure 7-figure supplement 1C source data 1). Rather, MEME-ChIP TF motif analysis suggests a striking change in the TFs that Tle1-4 associated with when *Sp5/8* were mutated. While Tcf/Lef motifs were not associated with Tle1-4 peaks in CHIR-stimulated WT EBs, Tbx, Tcf/Lef and Sp/Klf motifs, amongst others, were significantly associated with Tle peaks in *Sp5/8 dko* EBs (Figure 7-figure supplement 2A, B). These findings demonstrate that the reduced expression of Wnt target genes in *Sp5/8 dko* cells correlates with an increased association of Tle with WREs. Gene ontology and KEGG pathway analysis of the 266 genes that displayed both reduced Tcf7 peak intensities and increased Tle peaks in *Sp5/8 dko* cells compared to WT cells confirmed that these genes are associated with Wnt signaling, stem cells, gastrulation and AP patterning and include *Cyp26a1, Msgn1*, and *Sp5* (Figure 7B, Figure 7-figure supplement 2C, D, Supplementary file 6).

Detailed assessments of ChIP genome tracks at major regulators of axial progenitors for which there are well-characterized WREs, specifically *Tbxt, Wnt3a, Cdx2, Msgn1, Fgf8* and *Tbx6* (Amblard et al., 2024; Chalamalasetty et al., 2011; Nowotschin et al., 2012; Yamaguchi et al., 1999) showed that these WREs are bound by multiple Wnt/β-catenin transcriptional regulators i.e. Sp5, β-catenin, Lef1, Tcf7 and Tle (Figure 7C-H). Consistent with the genome-wide analysis, the Tcf7 peaks were reduced in intensity at 5 of 6 of these NMP genes in *Sp5/8 dko* EBs (Figure 7D-H). The loss of Tcf7 binding in *Sp5/8 dko* mutants was most notable at the *Wnt3a-DE* and the *Msgn1* promoter, and this correlated well with the reduced transcript levels (Figure 7D, E, top rows). ChIP-qPCR validated the ChIP-seq results for *Wnt3a* and *Msgn1.* Interestingly, despite our inability to observe differential binding of Tcf7 at the *Tbxt* promoter by ChIP-seq, ChIP-qPCR detected reduced Tcf7 binding in *Sp5/8 dko* EBs (Figure 7C). In contrast to the reduced Tcf7 binding, elevated Tle peaks were observed at the corresponding WRE for 5 of 6 genes in *Sp5/8* mutant cells (Figure 7C-E, G, H). The increase in Tle binding was apparently not mediated by Tcf7l1 since no change in Tcf7l1 binding was observed in *Sp5/8* mutants by ChIP-qPCR (Figure 7-figure supplement 3). Together, these results suggest that Sp5 enhances Tcf7, and reduces Tle, occupancy at the WREs of active Wnt target genes.

## Discussion

### Sp5/8 are integral components of the Wnt3a-Tbxt feedback loop in axial progenitors

Our findings demonstrate that Sp TFs are essential regulators of axial progenitor homeostasis, functioning to control morphogen signaling and TF networks within the NMP niche. Tissue-specific deletion of *Sp5* and *Sp8* in the PS resulted in a severe axial truncation due to the depletion of trunk axial progenitors. Notably, the anterior positioning of the axial truncation in *Sp5/8* double mutants, compared to *Sp8* single mutants which display a more posteriorly positioned tail truncation (Bell et al., 2003; Harrison et al., 2000), suggests a dose-dependent requirement for Sp5/8 in axial elongation that closely resembles previously described dose-dependent phenotypes in the Wnt pathway, for ex. *Wnt3a;8a*, *Wnt3a;vestigial tail, Tcf1;Lef1, Cdx1;2*, and *Cdx2; Tbxt* compound mutants (Chawengsaksophak et al., 2004; Chawengsaksophak et al., 1997; Galceran et al., 1999; Greco et al., 1996; MacMurray and Shin, 1988; Savory et al., 2009; Stott et al., 1993; van den Akker et al., 2002; van Nes et al., 2006; Young et al., 2009). Genetic interactions between *Sp5* and *Wnt3a* (Dunty et al., 2014), *Sp5* and *Tbxt* (Harrison et al., 2000), and *Tbxt* and *Cdx2* (Amin et al., 2016) firmly place *Sp5* and, by extension, *Sp8,* in this genetic *Wnt3a*-*Tcf1/Lef1*-*Tbxt*-*Cdx2* axis. Increasing activity of this canonical Wnt/β-catenin pathway is necessary for the emergence of increasingly posterior progenitors.

Axial progenitors must persist in the CLE and PS and, later, in the tailbud for at least 5 days of embryonic development (E7.5-12.5) to generate the entire trunk and tail (Cambray and Wilson, 2002, 2007). As axial development progresses in an anterior-to-posterior direction, early disruptions in progenitor maintenance result in anterior truncations, while later disruptions cause posterior truncations. The anterior truncation observed in *Sp5/8* double mutants aligns with an early requirement in trunk progenitors while the *Sp8* single mutant phenotype suggests a later requirement in tail progenitors. A plausible interpretation of the correlation between axial position and *Wnt3a* or *Sp5/8* gene dosage is that increasing Wnt activity is required to sustain axial progenitor self-renewal over time. In other words, Wnt/Sp signaling regulates the orderly timing of posterior progenitor emergence. The higher the Wnt/Sp signal, the later posterior progenitors emerge from the PS. This is supported by GOF studies in which overexpression of Sp8 in axial progenitors led to their retention in the caudal progenitor zone. We suggest that elevated levels of Sp sustain the *Wnt3a-Tbxt-Cdx* feedback loop for longer, thereby maintaining the self-renewal of increasingly posteriorized progenitors, possibly through Cdx and the maintenance of the Hox clock (Deschamps and Duboule, 2017).

The identification of Sp5 and Sp8 as both downstream targets and upstream regulators of *Wnt3a*, *Tbxt* and *Cdx2* strongly implicates their participation in an autoregulatory loop essential for axial progenitor maintenance. Previous work identified *Tbxt* as a direct target of Wnt3a/β-catenin signaling (Arnold et al., 2000; Yamaguchi et al., 1999), but the mechanisms governing *Wnt3a* activation by Tbxt remained elusive. Our data now reveal that Tbxt, Sp5, Sp8, Cdx2, β-catenin and Tcf7 converge on a conserved enhancer downstream of *Wnt3a* to regulate enhancer accessibility and activity. Single-cell ATAC-seq analysis in *Tbxt* mutants confirmed a crucial role for Tbxt in maintaining chromatin accessibility at this enhancer, reflecting a broader role of Tbxt in regulating the axial progenitor genetic program (Koch et al., 2017). Examination of WREs at *Wnt3a*, *Fgf4, Fgf8, Sp5*, *Sp8*, *Lef1, Tbxt, Cdx1, 2* and 4 reveals the regulatory logic that Tbxt and associated TFs use to activate these axial progenitor genes - they are bound by Tbxt, Tcf7, and either Cdx2, Sp5 or Sp8 (this work; (Amin et al., 2016; Kennedy et al., 2016; Koch et al., 2017; Savory et al., 2009; Zhu and Lohnes, 2022). We propose that reciprocal interactions among Sp5, Sp8, Tcf7, Tbxt and Cdx TFs constitute a robust transcriptional module that collectively sustains *Wnt3a*, *Fgf4* and *Fgf8* expression in axial progenitors through positive feedback.

### Sp5/8 and the NMP niche

A stem cell niche is defined as a specialized microenvironment that maintains progenitor self-renewal. The niche contains support cells that provide a local source of signals that regulate stem cell homeostasis. In the human adult intestinal stem cell (ISC) niche, ISCs are maintained by Wnt3a secreted by neighboring epithelial Paneth cells and underlying mesenchymal stromal cells (Clevers et al., 2014; Santos et al., 2018). By analogy, the self-renewal of embryonic NMPs relies on Wnt3a, however, the cellular composition of the NMP niche is inadequately defined. Our results suggest that the PS itself acts as a critical signaling center in the NMP niche. Axial progenitors residing in the CLE are likely regulated by both autocrine and paracrine signaling as *Wnt3a, Wnt8a, Fgf4, Fgf8* and *Fgf17* are expressed in NMPs and, more broadly, throughout the PS (Cunningham et al., 2015; Dunty et al., 2008; Dunty et al., 2014; Gouti et al., 2017; Koch et al., 2017; Maruoka et al., 1998; Pijuan-Sala et al., 2019; Tran et al., 2024; Wahl et al., 2007). Together with the high levels of Wnt and Fgf emanating from the posteriorly positioned PS, low levels of RA secreted from anteriorly positioned somites define the signaling environment that maintains NMPs (Gouti et al., 2017). Reduced expression of the RA-metabolizing enzyme *Cyp26a1* in *Sp5/8 dko* axial progenitors indicate that Sp5 and Sp8 also regulate RA levels in the NMP niche (Figure 4-figure supplement 2). We conclude that Sp5/8 play a central role in the NMP niche to establish a high Wnt/Fgf, low RA signaling environment that ensures spatially coordinated neural and mesodermal differentiation.

In addition to functioning as the Left-Right (LR) organizer, the node has also been implicated as a niche component that sustains NMPs through paracrine signals (Wymeersch et al., 2019). The node largely consists of ciliated notochord progenitors and lies ventral to, and in direct contact with, NMPs throughout axial elongation. Localized ablation of ventral node cells at the NSB at E8.5 perturbed axis elongation, leading to speculation that the node provides signals that sustain NMPs. The observed expression of *Sp5/8* in the node raises the intriguing possibility of their involvement in node signaling. Indeed, reduced nodal cilia have been observed in *Sp5/8 dko* embryos, leading to altered LR patterning and organ laterality defects (Liang et al., 2025). Cilia are signal-processing organelles that sample chemical and mechanical cues in the cellular signaling environment and translate them into intracellular responses. Although the requirement for *Sp5/8* in node cilia formation is intriguing, a role for node cilia in NMP maintenance has not been established. Node cilia project from the apical surface of node cells, away from NMPs which are in contact with the basal surface of node cells and separated by ECM. Nevertheless, we suggest that the NMP niche is composed of two neighboring but spatially distinct support cell populations, the PS, lying caudal to NMPs, and the node lying immediately ventral.

### Sp5/8 and Wnt target gene regulation

Given the crucial roles that the Wnt/β-catenin pathway plays in the regulation of gene expression in embryonic, adult and cancer stem cells, it is of fundamental importance to understand how the pathway regulates transcription. β-catenin and the four Tcf/Lef TFs, Tcf7, Lef1, Tcf7l1 and Tcf7l2, are the primary transcriptional effectors of canonical Wnt signaling. Vertebrate Tcfs function predominantly as dedicated activators (Lef1), repressors (Tcf7l1) or both (Tcf7 and Tcf7l2) depending on the cellular context (reviewed in (Bou-Rouphael and Durand, 2021; Guo et al., 2021; Ramakrishnan et al., 2018). Tcf repressor activity correlates with their affinity for Tle co-repressors as the dedicated repressor Tcf7l1 binds strongly to Tle while the activator Lef1 does not (Chodaparambil et al., 2014). Notably, in pluripotent ESCs stimulated with Wnt3a, β-catenin binding to Tcf7l1-Tle triggers its eviction from chromatin and subsequent degradation (Shy et al., 2013). In contrast, β-catenin binding to Tcf7 promotes the formation of a chromatin-bound transcriptional activation complex (Yi et al., 2011). The molecular mechanisms that determine these distinct outcomes of β-catenin binding remain poorly understood. Nevertheless, the β-catenin-dependent replacement of repressive Tcf7l1-Tle with activating Tcf7-β-catenin at WREs is consistent with an exchange of Tcfs driving transcription. It should be noted, however, that Tcf exchange could operate in parallel with a Tcf7 switch mechanism whereupon both Tcf7-Tle and Tcf7l1-Tle repress Wnt targets in cells that coexpress Tcf7 and Tcf7l1.

Our ChIP-seq studies investigating the genomic distribution of Sp5/8 show that they are bound to many of the same WREs as the primary Wnt transcriptional effectors β-catenin, Tcf7, and Lef1. These WREs are also bound by Tbxt and Cdx TFs which, like Sp5/8, are regulated by, and required for, transcriptional responses to Wnt. Given that Tcf7 and Lef1 proteins can physically interact with Sp5/8 (Kennedy et al., 2016) and Cdx1 (Beland et al., 2004; Ramakrishnan et al., 2021), we suggest that Tbxt, Cdx1/2 and Sp5/8 are context-dependent secondary Wnt transcriptional effectors that cooperate with Tcf/β-catenin at WREs to establish primitive streak-specific transcriptional programs.

Mechanistically speaking, how do secondary effectors like Sp5/8 regulate the principal effectors of Wnt signaling? We have presented evidence that Sp5/8 promote Tcf7 occupancy at WREs of actively transcribed genes. Tcf7 peaks were frequently reduced at WREs in the absence of Sp5/8, but Tcf7 nevertheless remained detectable and bound to canonical Tcf motifs (Figure 7-figure supplement 2A) indicating that the presence of Sp5/8 does not alter Tcf7 DNA binding specificity. Interestingly, robust binding of Tle co-repressors was observed at many sites of reduced Tcf7 binding suggesting that repressive Tcf-Tle complexes were maintained at WREs in *Sp5/8* mutants despite the presence of CHIR. ChIP-qPCR analysis at select WREs (*Tbxt, Sp5, Msgn1* and *Wnt3a*) did not detect enrichment of Tcf7l1 (Figure 7-figure supplement 3), indicating that Tcf7 or Tcf7l2 is likely responsible for Tle recruitment to WREs. We conclude that Sp5/8 influence how Tcf complexes respond to Wnt activation and suggest that Sp5/8 act cooperatively to enhance the binding of Tcf7-β-catenin to WREs. It is worth considering that other non-Tcf TFs could also participate in Tle recruitment since motifs for Tbx and Lhx, amongst others, were enriched at Tle peaks in *Sp5*/8 mutants.

A role for Sp TFs in Wnt/β-catenin signaling appears to be evolutionarily conserved (Dailey et al., 2017; Dunty et al., 2014; Gautam et al., 2024; Mohanty and Lekven, 2025; Tewari et al., 2019; Weidinger et al., 2005). Like Tcf/Lef factors, Sp proteins can act as either activators or repressors, suggesting that their activity depends on interactions with cofactors. In mouse, Sp5 and Sp8 promote *Wnt3a* transcription in NMPs (this work; (Kennedy et al., 2016). Sp8 also activates *Fgf8* in the limb (Kawakami et al., 2004), and represses *Emx2* in the forebrain (Zembrzycki et al., 2007). Similar context-dependent activities are seen in other systems. For example, in the regenerating acoel, Hofstenia miamia, *Sp5* represses Wnt-regulated trunk genes such as *Tbxt* and *Hox* in the tail (Tewari et al., 2019). In *Hydra,* Sp5 represses *Wnt3* to prevent ectopic head formation (Vogg et al., 2019), while in differentiating human PSCs, Sp5 suppresses a Wnt-dependent transcriptional program that produces cardiac mesoderm and definitive endoderm amongst others (Huggins et al., 2017). Although these latter findings differ from the gastrulation data presented here, they likely reflect a different developmental context, as our work examines later-arising trunk NMPs, whereas Huggins et al. studied cells that differentiate during an earlier stage of gastrulation. Together, these studies support the view that Sp factors are bifunctional, context-dependent, secondary Wnt transcriptional effectors.

## Supplementary Figures

**Figure 2-figure supplement 1.**
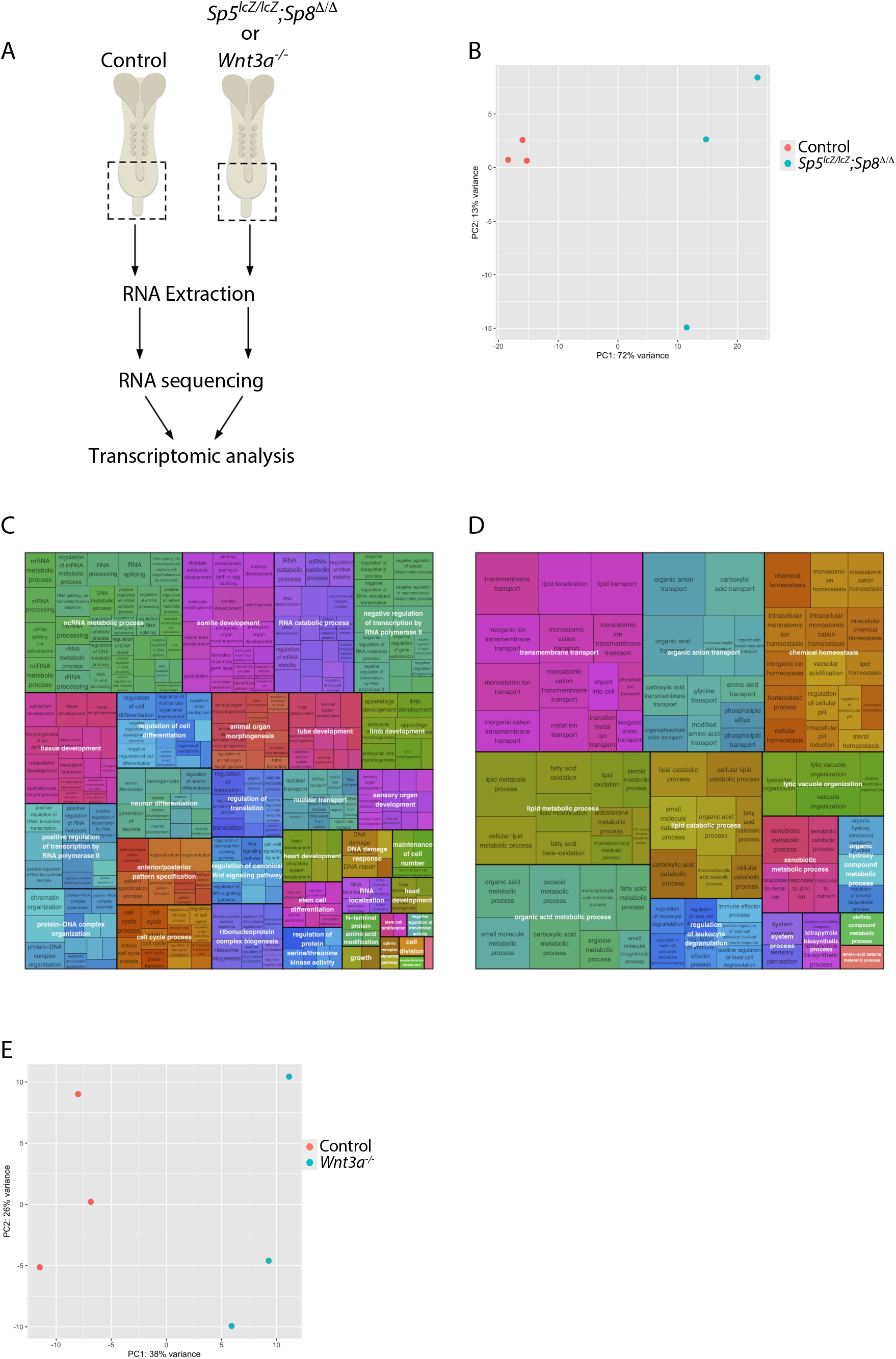
Gene Set Enrichment Analysis (GSEA) of differentially expressed genes in *Sp5/8 dko* bulk-RNA seq. Related to Figure 2. A. Schematic representation of E8.5 control and *Sp5/8 dko* or *Wnt3a^-/-^* mutants, with the dashed box illustrating the posterior termini dissected for RNA extraction and bulk-RNA sequencing (n=3 for controls and *Sp5/8 dko* mutants). B. Principal component analysis of control (n=3) and *Sp5/8 dko* (n=3) samples used for bulk-RNA seq analysis. C-D. Treemap visualization of significantly enriched GO terms identified through GSEA analysis of *Sp5/8 dko* bulk RNA-seq differentially expressed downregulated genes (E) and upregulated genes (F). Each rectangle denotes a GO term, with similar terms grouped by color to reflect semantic similarity. Rectangle areas are proportional to adjusted p-values (−log10 transformed). E. Principal component analysis of control (n=3) and *Wnt3a^-/-^* (n=3) samples used for bulk-RNA seq analysis.

**Figure 3-figure supplement 1.**
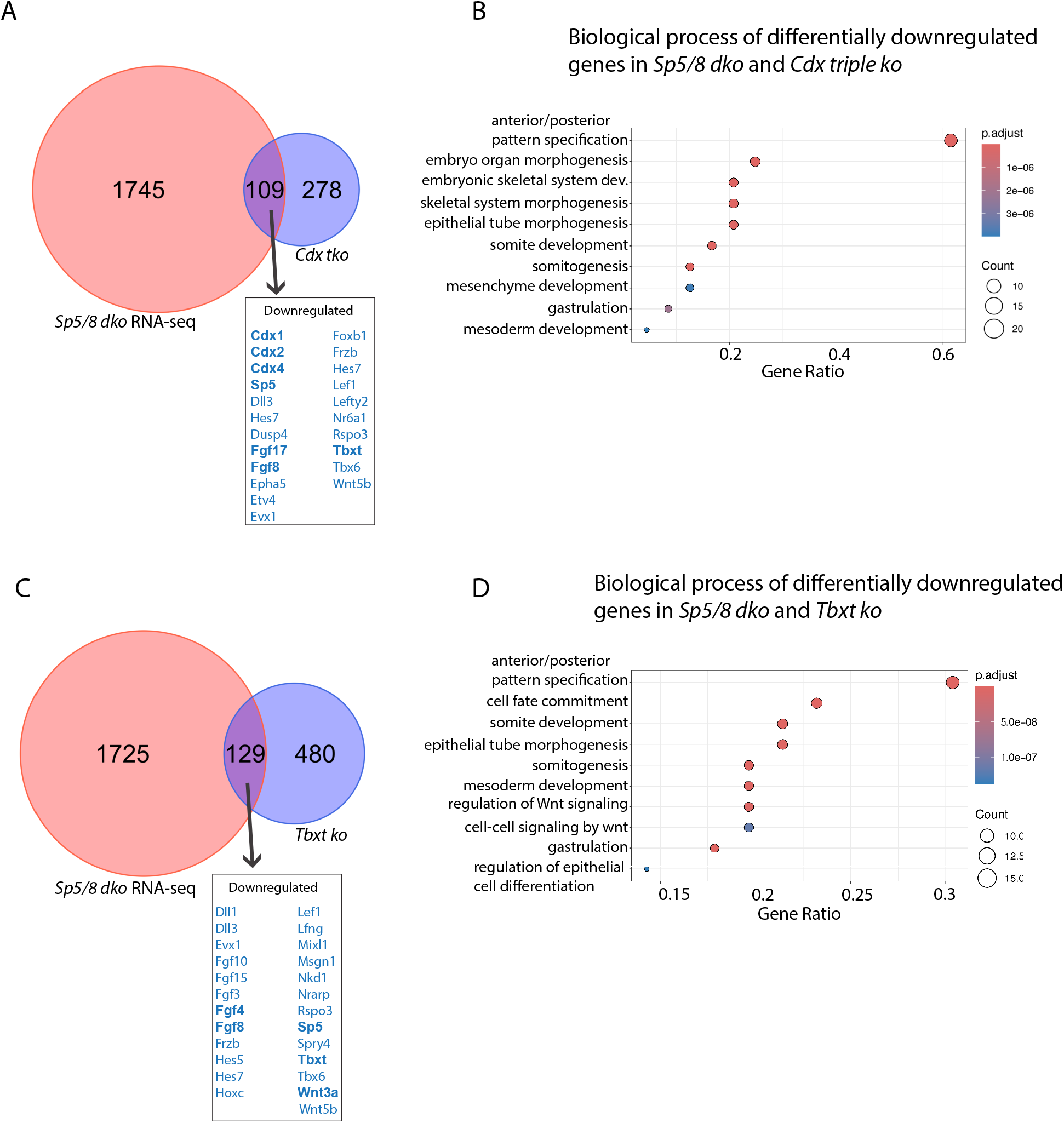
Comparisons of *Sp5/8 dko* differentially expressed genes to Cdx*- tko* and Tbxt *ko*. Related to Figure 3. A. Venn diagram showing the overlap of DEGs between the *Sp5/8 dko* and *Cdx tko* (Amin et al., 2016). Select downregulated genes are featured, NMP-relevant genes are highlighted in bold. B. GO biological processes terms of downregulated genes shared between *Sp5/8 dko* and *Cdx tko* datasets. C. Venn diagram showing the overlap of DEGs between *Sp5/8 dko* and *Tbxt ko* (Koch et al., 2017). Key down regulated genes are highlighted. D. GO biological processes terms of downregulated genes shared between *Sp5/8 dko* and *Tbxt* ko datasets.

**Figure 4-figure supplement 1.**
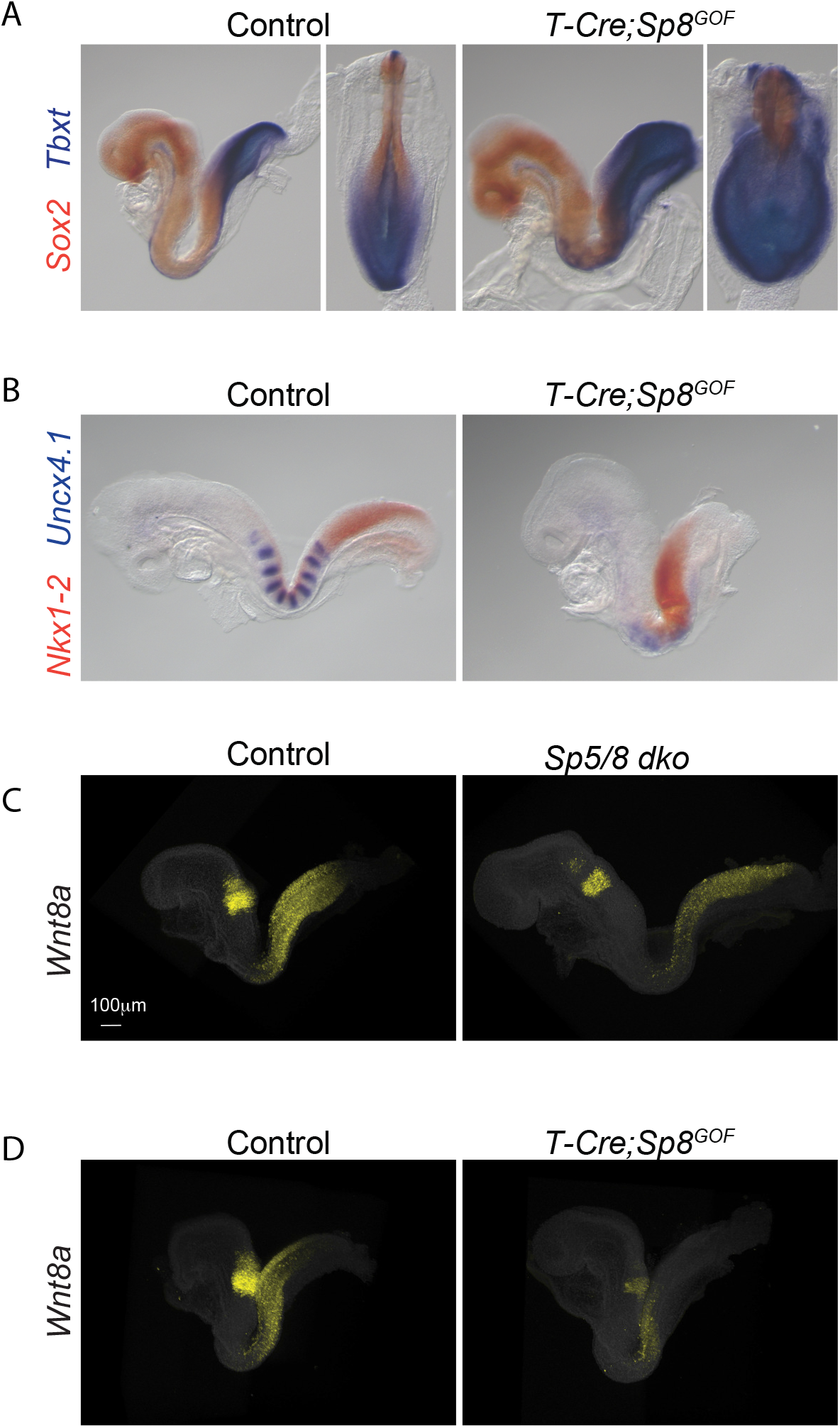
RNA in situ expression of NMP genes in *T-Cre-Sp8^GOF^* and *Sp5/8 dko* embryos. Related to Figure 4. A-B. Two color whole mount in situ hybridization of E8.5 embryos for *Sox2* (orange) and *Tbxt* (purple) (A) and *Nkx1-2* (orange) and *Uncx4.1*(purple) (B) in control and *T-Cre; Sp8^GOF^* embryos. Lateral and dorsal views are shown A and lateral view in B. C-D. Lateral view of whole mount fluorescent in situ hybridization analysis for transcripts of *Wnt8a* (yellow) in *Sp5/8 dko* embryos (C) and *T-Cre; Sp8^GOF^* embryos at E8.5 (D).

**Figure 4-figure supplement 2.**
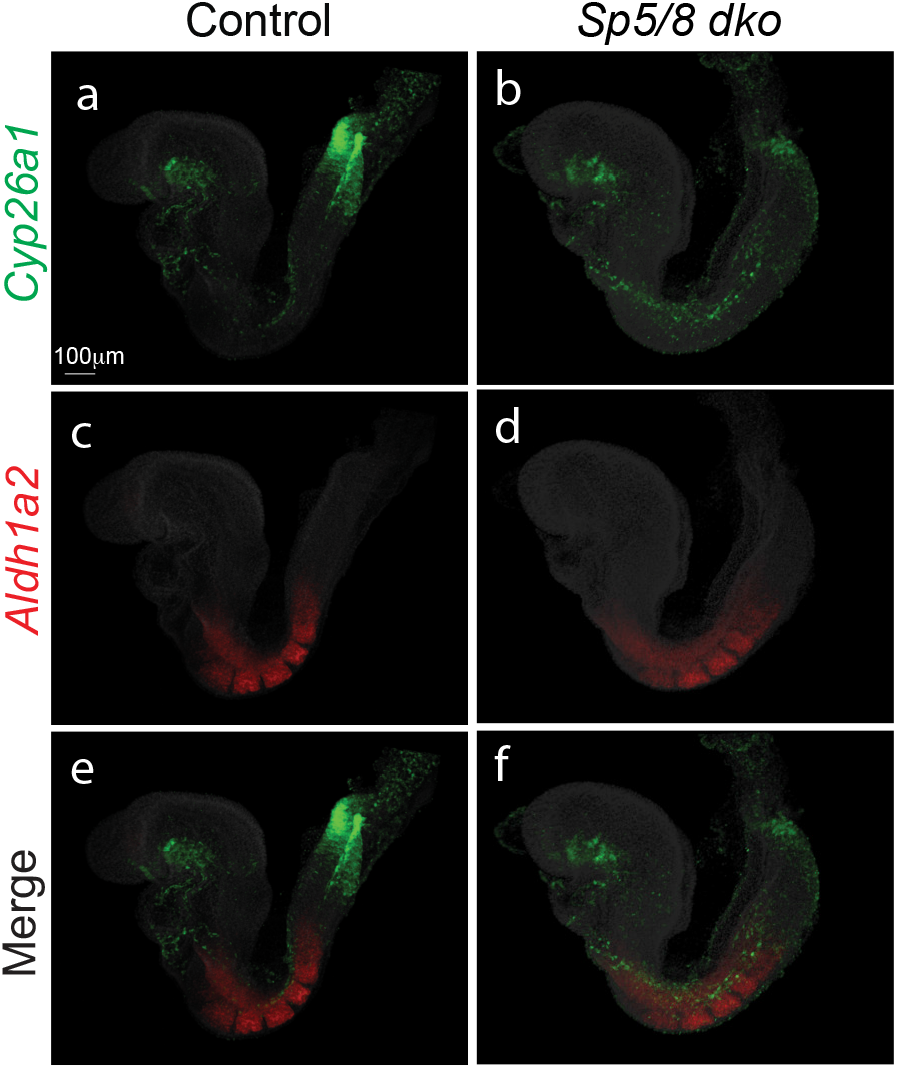
Whole mount fluorescent in situ hybridization analysis of RA pathway genes. Related to Figure 4. Lateral view of embryos processed for *Cyp26a1* (green; a, b, e, f) and *Aldh1a2* (red, c, d, e, f) in E8.5 control and *Sp5/8 dko* embryos.

**Figure 5-figure supplement 1.**
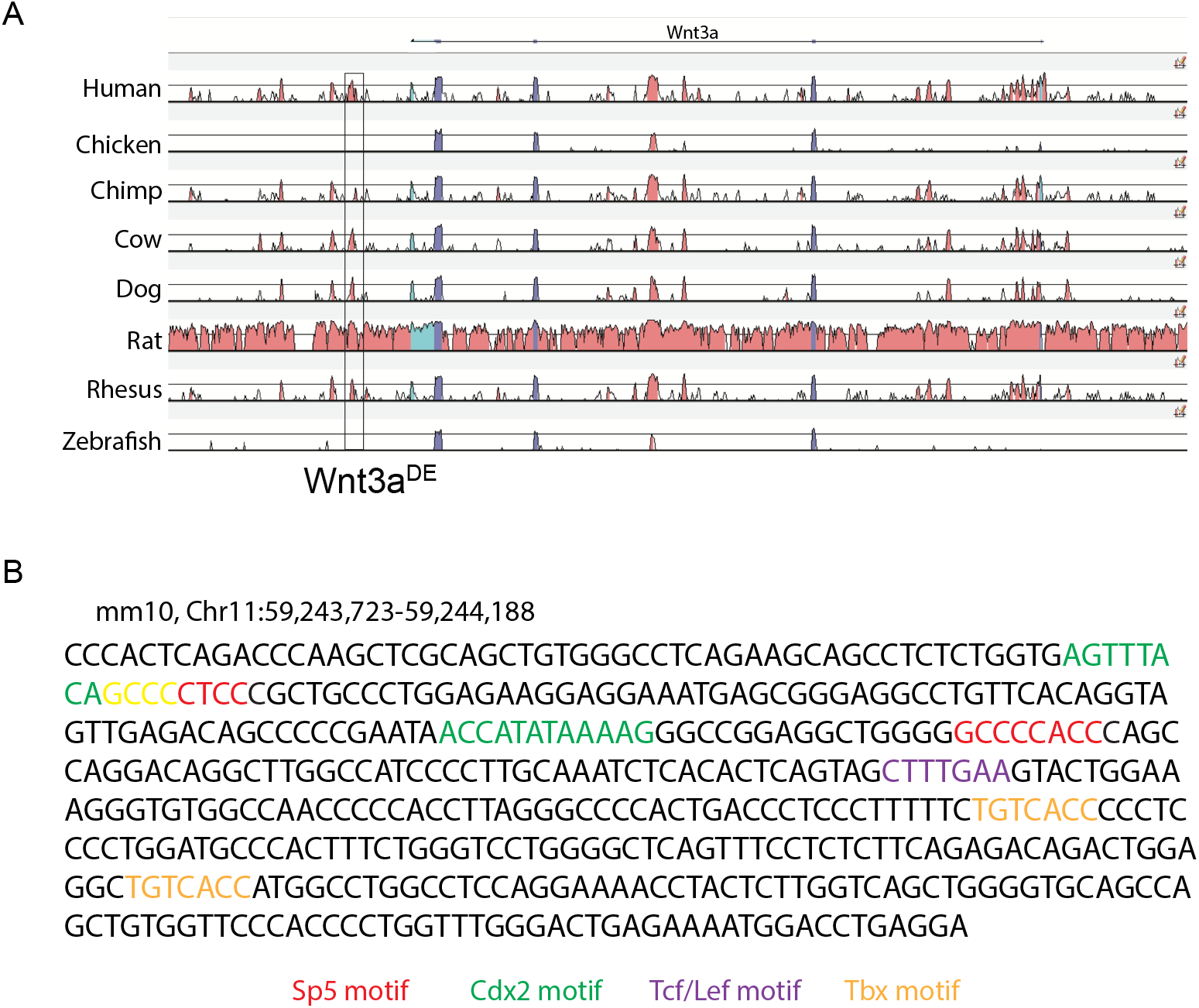
Characterization of *Wnt3a^DE^* using comparative genomics. Related to Figure 5. A. Comparative genomics analysis of the mouse Wnt3a locus using Vista tools (Frazer et al., 2004). The conserved *Wnt3a^DE^*is boxed. B. Motif annotations from CIIIDER annotated against the 466 bp putative Wnt3a^DE^ sequence (Gearing et al., 2019). Wnt3a^DE^ is identified as −15696 intergenic peak (mm9; Chr11: 59057304-59057626 upstream of Arf1(see Supplementary file 1)

**Figure 6-figure supplement 1.**
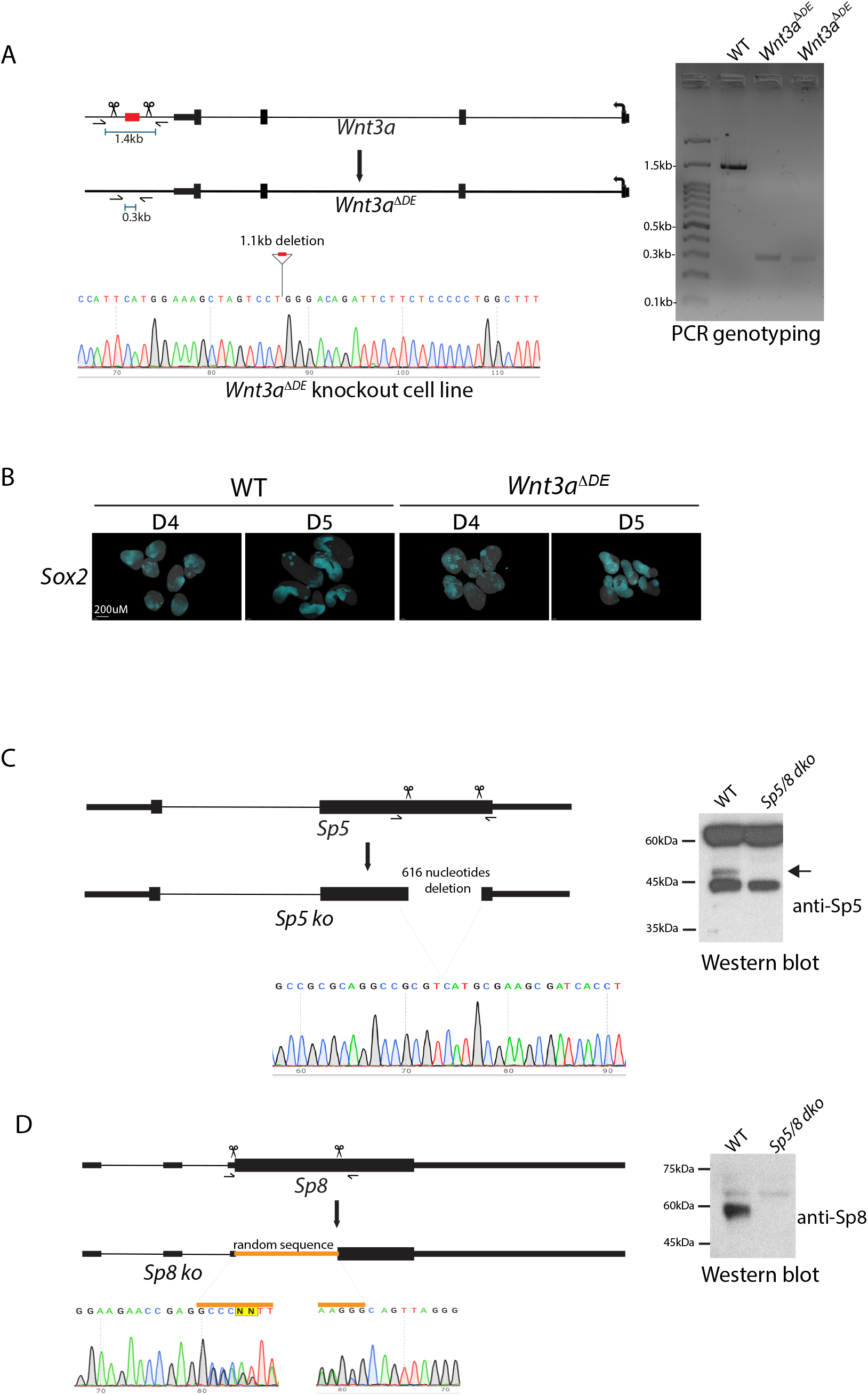
Generation of *Wnt3a^ΔDE^* and *Sp5/8 dko* ESCs using CRISpR/cas9 technology. Related to Figure 6. A. Schematic view of *Wnt3a^ΔDE^* ko ES cell generation using CRISpR-Cas9 mediated deletion. Two guides (see Supplementary file 7) were designed to delete 1.1kb encompassing the 0.47kb *Wnt3a^DE^*. Genotyping oligos were designed to amplify a 1.4kb fragment in wildtype and 0.3kb fragment in *Wnt3a^ΔDE^* ko ESCs. PCR genotyping gel shows deletion of *Wnt3a^DE^* in two different clones. B. HCR analysis of *Sox2* (cyan) expression in WT and *Wnt3a^ΔDE^* gastruloids one (D4) and two (D5) days after CHIR treatment, respectively. Dapi, grey. C-D. Generation of *Sp5/8 dko* ES cells. C. Schematic of *Sp5 ko* using CRISpR-Cas9 mediated deletion in Exon-2 using two guides designed to delete 616 bases encoding the Zn-finger domains of Sp5. ES cells were differentiated as EBs and treated with bFgf+CHIR for 24h. Sp5 protein was not detected in D3 cell extracts. D. Schematic view of *Sp8 ko* using CRISpR-Cas9 mediated deletion in Exon-3 using two guides designed to delete the coding region. Random sequence was inserted by NHEJ repair and these ES cells failed to express Sp8 protein in D3 EBs.

**Figure 7-figure supplement 1.**
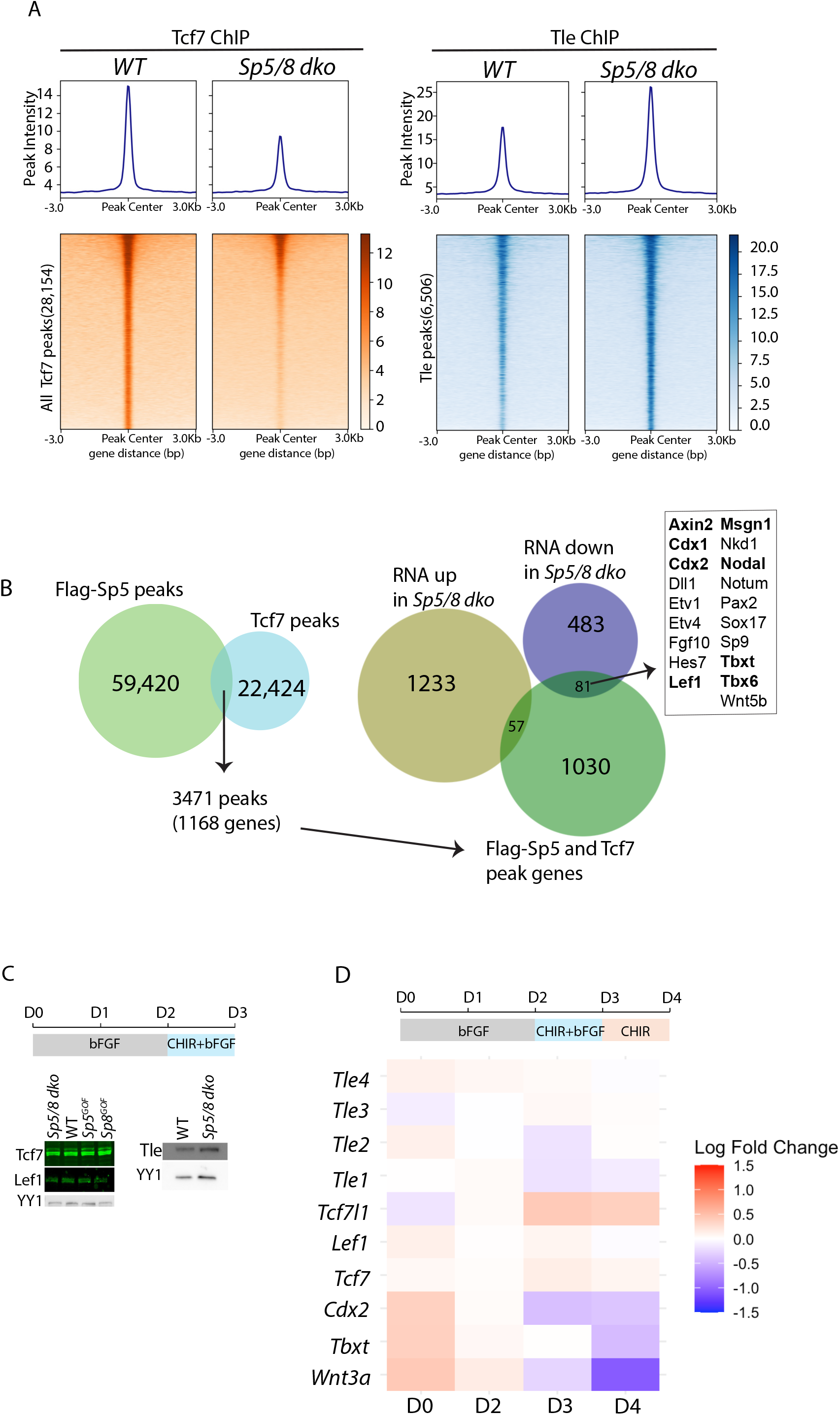
Comparative analyses of Flag-Sp5, Tcf7, and Tle ChIP-seq data sets. Related to Figure 7. A. Metaplots and heat maps depicting 28,154 Tcf7 ChIP-seq peaks and 6,506 Tle peaks in WT and *Sp5/8 dko* EBs. B. (Left)Venn diagram illustrating the overlap between Flag-Sp5 and Tcf7 ChIP-seq peak sets. A total of 3,471 shared putative WREs, corresponding to 1168 unique genes, were identified. (Right) Venn diagram showing the intersection of the 1,168 genes associated with shared Flag-Sp5 and Tcf7 peaks and genes differentially expressed (up- or downregulated) in *Sp5/8 dko* bulk-RNA-seq. C. (Left) Western blot analysis depicting Tcf7 and Lef1 protein expression in NMPs differentiated in vitro from WT, *Sp5/8 dko*, *Sp5^GOF^* and *Sp8^GOF^* EBs. (Right) Tle protein expression in WT and *Sp5/8 dko* NMPs differentiated as EBs. YY1, loading control. D. Quantitative PCR time course showing log fold change of RNA expression of *Tle1-4, Tcf7l1, Tcf7, Cdx2, Tbxt,* and *Wnt3a* in *Sp5/8 dko* versus WT EBs. RNA expression was normalized to *Gapdh* for all genes.

**Figure 7-figure supplement 2.**
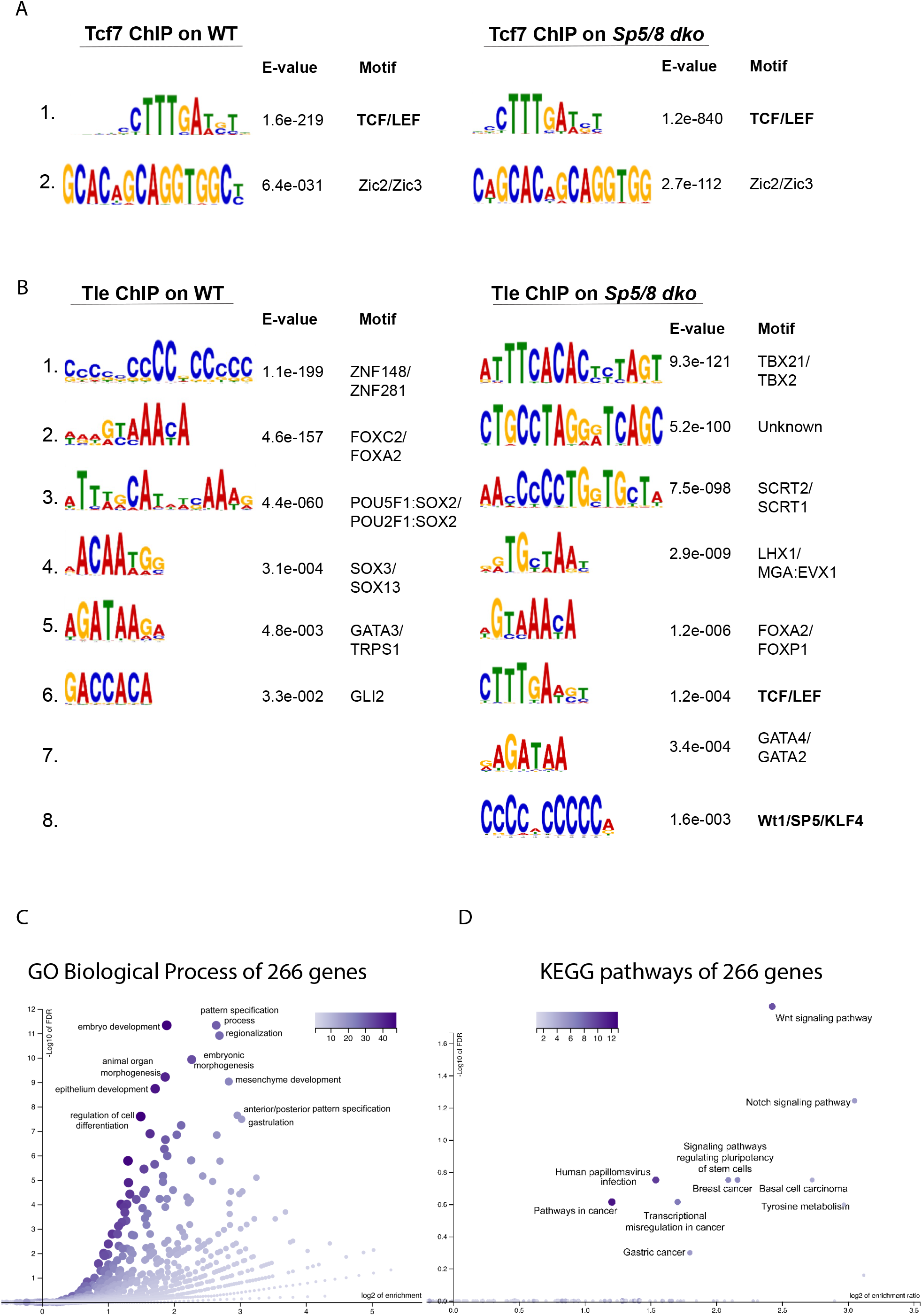
Motif and Over-representation analyses of F-Sp5, Tcf7, and Tle ChIP-seq datasets. Related to Figure 7. A-B. Meme-ChIP motif analysis of peak sequences from Tcf7 (A) and Tle (B) ChIP-seq data in WT and *Sp5/8 dko* cells. In Tcf7 ChIP-seq, the Tcf/Lef motif was the most significantly enriched motif in both WT and *Sp5/8 dko.* In Tle ChIP-seq, Znf motif and Tbx motif were most enriched in WT and *Sp5/8 dko,* respectively, with additional enrichment of Tcf/Lef and Sp/KLF motifs in *Sp5/8 dko* only. C-D. Over-representation analysis (ORA) using WEB-based GEne SeT Analysis Toolkit on 266 overlapping genes corresponding to 286 F-Sp5 and Tcf7 overlapping WREs (as shown in Figure 7B). Enrichment results are shown for (A) Gene Ontology (GO) Biological Process terms and (B) KEGG pathways.

**Figure 7-figure supplement 3.**
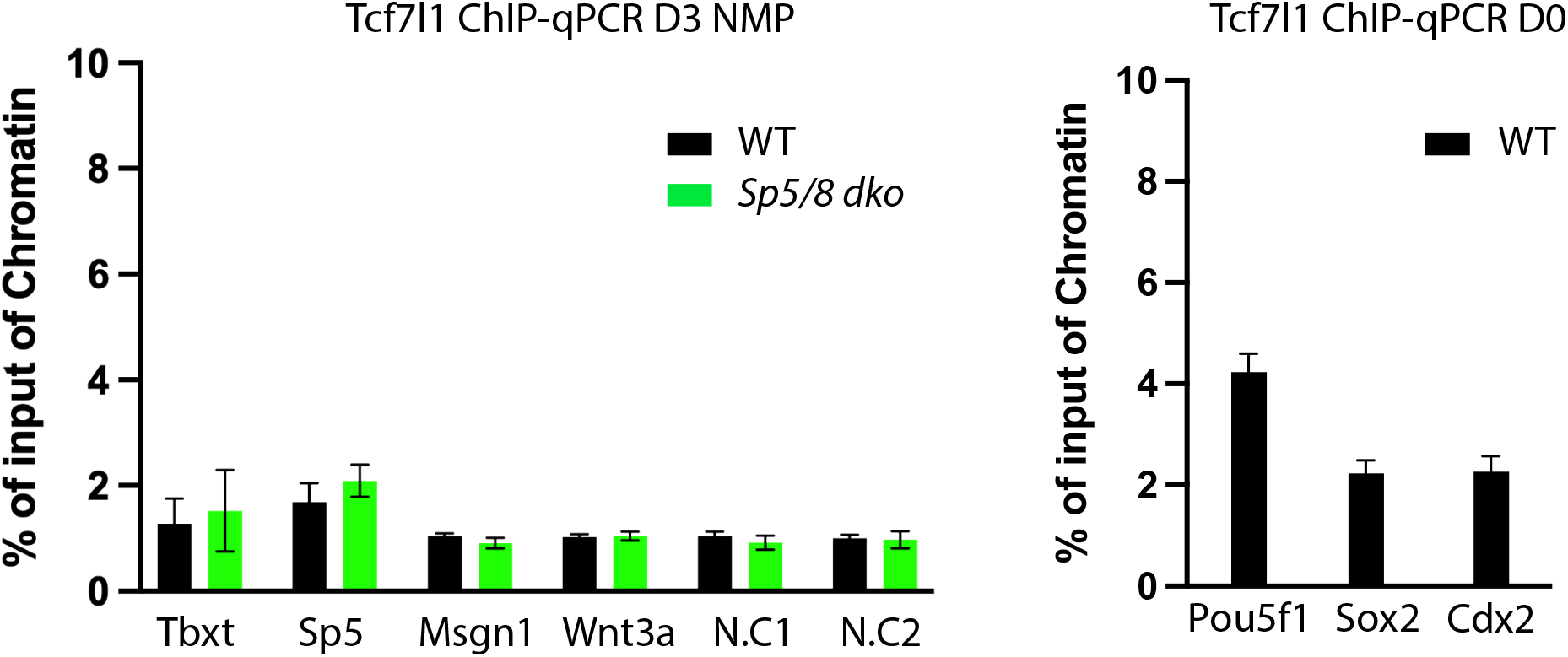
Tcf7l1 ChIP-qPCR at WREs in WT and *Sp5/8 dko* day 3 EBs. Related to Figure 7. ChIP-qPCR analysis of Tcf7l1 occupancy at regulatory regions associated with *Tbxt, Sp5, Msgn1*, and *Wnt3a* in day 3 EBs derived from WT (black) and *Sp5/8 dko* (green) cells. Regulatory regions associated with *Pou5f1, Sox2* and *Cdx2* in pluripotent ESCs served as positive controls for the Tcf7l1 antibody. Enrichment is plotted as percent of input chromatin. Bars depict the mean, and error bars denote standard deviation. N.C1 and N.C2 indicate negative-control regions designed within the *Tbxt* locus (Supplementary file 7).

Supplementary file 1

Overlap of *Sp5/8* bulk-RNA seq with Sp5 ChIP-seq (GSE73084) data. Related to Figure 2C Supplementary file 2

Overlap of *Sp5/8 dko* and *Wnt3a ko* bulk RNA seq. Worksheets contain both unique, shared, and shared downregulated gene sets. Related to Figure 2D

Supplementary file 3

Comparative transcriptome analysis of *Sp5/8 dko, Wnt3a ko, Cdx tko, Tbxt ko* data sets. Related to Figure 3A and Figure 3-figure supplement 1.

Supplementary file 4

Comparative ChIP-seq analysis of Flag-Sp5, Flag-Sp8, Cdx2 and Tbxt datasets. Related to Figure 3B.

Supplementary file 5

List of all ChIP-seq peaks for Flag-Sp5, Tcf7 and Tle conditions. Related to Figure 7, and Figure 7-figure supplement 1.

Supplementary file 6

Summary of all ChIP-seq data comparisons. Related to Figure 7, and Figure 7-figure supplement 1.

Supplementary file 7

Details of gRNAs, sequencing primers for characterizing knockout cell lines, qPCR oligos, ChIP-qPCR oligos, GEO accession numbers and antibodies for Immunohistochemistry, ChIP-seq, ChIP-qPCR, Immunoprecipitations, and western blots are included.

Source code 1

R and Shell scripts for bulk-RNAseq analysis of Sp5/8 dko, Wnt3a ko and control embryos.

Figure 6-figure supplement 1A source data 1

Uncropped raw agarose images of Figure 6-figure supplement 1A source data 1

Figure 6-figure supplement 1A source data 2

Cropped raw agarose images of Figure 6-figure supplement 1A source data 2

Figure 6-figure supplement 1C source data 1

Uncropped raw blot images of Figure 6-figure supplement 1C source data 1

Figure 6-figure supplement 1C source data 2

Cropped raw blot images of Figure 6-figure supplement 1C source data 2

Figure 6-figure supplement 1D source data 1

Uncropped raw blot images of Figure 6-figure supplement 1D source data 1

Figure 6-figure supplement 1D source data 2

Cropped raw blot images of Figure 6-figure supplement 1D source data 2

Figure 7-figure supplement 1C source data 1

Uncropped raw blot images of Figure 7-figure supplement 1C source data 1

Figure 7-figure supplement 1C source data 2

Cropped raw blot images of Figure 7-figure supplement 1C source data 2

## Materials and Methods

### Mouse strains and skeletal preparation

*Sp5^lacZ/lacZ^; Sp8^+/Δ^*, *T-Cre, Sp8^GOF^ (TetO-Sp8-IRES-EGFP), Rosa26-rtTA-IRES-EGFP (*B6.Cg-*Gt(ROSA)26Sor^tm1(rtTA,EGFP)Nagy^*/J; RRID: IMSR_JAX:005670) mice were described previously (Dunty et al., 2014; Kennedy et al., 2016; Perantoni et al., 2005). *Sp8^GOF^* mice were crossed with *Rosa26-flox-rtTA* mice to enable temporal and conditional expression of Sp8 using the *T-Cre* line. For temporal expression, pregnant females were provided with Dox chow (200mg/kg) (Bio-Serv, S3888) and Dox water (1.6mg/ml wt/vol Dox) in 5% (wt/vol) sucrose from embryonic day 7.5 (E7.5) and were harvested at embryonic day 8.5 (E8.5). All mouse experiments in this study were performed in strict accordance with the animal protocol (Maintenance and characterization of mutant mouse strains, #26-408) approved by the NCI-Frederick Animal Care and Use Committee at the National Institutes of Health. Skeletal preparations of E18.5 fetuses were done as described previously (Hogan and Hogan, 1994).

### Whole mount in situ hybridization

For colorimetric whole mount in situ hybridization (WISH) experiments, DIG-labeled probes were synthesized, and WISH was conducted as described (Dunty et al., 2008) using a minimum of three embryos per genotype. For multiplexing whole mount Fluorescent In Situ Hybridization (FISH), split initiator hairpin probes (V 3.0) were custom-designed and synthesized by Molecular Instruments, Inc. Buffers and amplifiers conjugated to fluorophores were procured from Molecular Instruments, Inc, and FISH was carried out according to the manufacturer’s instructions (Choi et al., 2018). Embryos and gastruloids were harvested and fixed in 4% PFA overnight at 4^0^C and dehydrated in Methanol/PBST (PBS+0.1%Tween-20) series the following day. For HCR hybridization, embryos were rehydrated in a Methanol/PBST series, treated with Proteinase-K for 15 min followed by post-fixation in 4% PFA for 20 min at room temperature. After pre-incubation with probe hybridization buffer, samples were incubated with probe mix at 37^0^C for overnight. Samples were washed, and amplification was performed the next day in 60 pmol of each hairpin in 0.5 ml of amplification buffer overnight at room temperature. Next day, samples were washed in 5x SSCT buffer and stained in 0.5 ug/ml Dapi in 5x SSCT buffer overnight in 5x SSCT, then embedded on glass bottom dishes (MatTek or Greiner Bio-One GmbH) in 1% ultra-low melt agarose (Cambrex). Subsequently, samples were cleared at room temperature using Ce3D++ solution for 3-4 days before confocal imaging, as described (Anderson et al., 2020). Whole mount FISH was performed similarly for gastruloids (n=>5 per genotype), with the exception that gastruloids were not rotated on a shaker. All experiments were repeated at least 3 or more times.

### Whole mount immunohistochemistry

For Whole Mount Immunohistochemistry (WIHC), embryos (n=>3 per genotype) were fixed for 30 minutes in 4% paraformaldehyde at room temperature and subsequently washed in PBST (PBS+0.1% Tween-20). Permeabilization was achieved using 0.5% Triton-X 100, followed by blocking in 5% normal donkey serum in PBST for 1 hour. Embryos were then incubated in 100 μL of primary antibodies overnight at 4°C with shaking, followed by 5-6 PBST washes over 24 hours. Subsequently, embryos were incubated in Alexa Fluorophore-conjugated secondary antibodies (1:2000) at 4°C on a shaker, followed by another day of PBST washes containing 0.5 μg/mL DAPI. Finally, embryos were mounted as described above. Antibody information can be found in Supplementary file 7. All experiments were repeated at least 3 or more times.

### scRNA-seq and expression heat maps

UMAP projection plots for E8.25 embryo set for *Tbxt, Sox2, Sp5,* and *Sp8* were obtained from the interactive web portal in MouseGastrulationData2018 data (https://marionilab.cruk.cam.ac.uk/) (Pijuan-Sala et al., 2019). Heatmaps for expression of *Tbxt, Sox2, Sp5,* and *Sp8* from different treatment conditions in differentiating ESCs were extracted from multiplexed Barcodelet Single-Cell RNA-seq (Yeo et al., 2020).

### Bulk RNA-seq

For Bulk-RNA sequencing, embryos (n=3 per genotype) were collected at the 4-6 somite stage and bisected between somites (S)1 and S2. Posterior fragments were collected for RNA isolation, and anterior tissue was used for genotyping. Bulk RNA-seq paired end reads were processed using the RENEE v2.5.8 (Rna sEquencing aNalysis pipElinE) analysis pipeline (https://github.com/CCBR/renee) using default settings. Briefly, low-quality reads and adaptors were removed using Cutadapt v1.18 (RRID:SCR_011841) (Martin, 2011) before undergoing alignment to the GRCm38.p6 genome using STAR v2.7.6a (RRID:SCR_004463) in two-pass mode (Dobin et al., 2013). Post-alignment QC metrics were collected using Picard v 2.17.11 (RRID: SCR_006525) and RSeQC v2.6.4 (RRID:SCR_005275) while potential contamination was examined using the Kraken v2.0.8-beta (RRID:SCR_026838) and FastQ Screen v0.14.0 (RRID:SCR_000141) packages (Wang et al., 2012; Wingett and Andrews, 2018; Wood et al., 2019). Expected and isoform counts, according to the GENCODE vM25 (RRID:SCR_014966) primary assembly annotation reference, were collected using the RSEM v1.3.0 (RRID:SCR_000262) package. Differential expression analysis was performed using DESeq2 v1.42.0 (RRID:SCR_015687) in R v4.3.0 (Love et al., 2014; Soneson et al., 2015). Genes with less than 10 counts in at least 3 samples were filtered out. Genes were considered differentially expressed if they presented a log 2-fold change ≥0.5 or ≤-0.5 and an adjusted P-value ≤0.05 (Benjamini-Hochberg). The cluster Profiler v4.10.0 R/Bioconductor package (RRID:SCR_016884) was used to evaluate enrichment of Gene Ontology biological processes terms associated with lists of differentially expressed genes through Gene Set Enrichment Analysis (adjusted P-value <0.01) (Yu et al., 2012). We used the rrvgo package (RRID:SCR_026680) to reduce and interpret the list of associated GOs obtained from cluster Profiler (Sayols, 2023). We first defined two sets of GOs associated with either overexpressed or under expressed genes using their normalized enrichment scores. Each GO set was used to compute a similarity matrix considering the full set of mouse genes as background. We proceeded by computing the scores of each GO term, defined as the −log10 of their adjusted p-values, and used the similarity matrices, together with the list of scores, to obtain two lists of reduced terms (threshold of 0.7). Enhanced Volcano and Complex Heatmap and Venn Diagram packages were used for visualization throughout. For integrating published data sets, differentially expressed genes in the bulk-RNA seq for *T^2J/2J^*embryos (GSE93524)(Koch et al., 2017) and *Cdx* triple knockout embryos (GSE84899) (Amin et al., 2016) were used. Raw and processed sequencing files have been deposited at the GEO database (see Supplementary file 7 for accession numbers).

### Chromatin Immunoprecipitation and Sequencing (ChIP-seq)

ChIP-seq experiments analyzing Flag-Sp5 genome occupancy was described previously (Kennedy et al., 2016). ChIP-seq experiments for Tbxt, Flag-Sp5 and Flag-Sp8 were performed in A2lox.Cre ESCs (Iacovino et al., 2011), while Tle and Tcf7 ChIPs were conducted in R1 ESCs. At least two replicates per condition were used. ESCs were differentiated into NMPs as described (Gouti et al., 2014) in 2-dimensional cultures or embryoid bodies. Pluripotent ESCs were differentiated with bFgf (12ng/ml) for 2 days (D0-D2) followed by 3uM CHIR and 12ng/ml bFgf for 24 hours. For overexpression of inducible Flag-Sp5, 1ug/mL doxycycline (dox) was added from D2 to D3. Cells were harvested and fixed in 1% formaldehyde solution for 15min, quenched in 0.125M glycine and flash frozen. For ChIP-seq, nuclear extracts were incubated with antibodies as listed in Supplementary file 7. ChIP libraries were prepared and sequenced either on two separate NextSeq runs using V2 chemistry or on the NovaSeq Xplus 1.5B platform utilizing the IDT 2S Plus DNA library preparation kit (IDT) with single-end sequencing. All libraries demonstrated high sequencing quality, with over 94% of bases exceeding a Q30 quality score. Sequencing yield ranged from 42 to 67 million pass-filter reads per sample, indicating robust library preparation and sequencing performance. Downstream data processing and analysis were conducted using the CHAMPAGNE pipeline (DOI: <u>10.5281/zenodo.10516078</u>), a comprehensive and reproducible workflow developed for ChIP-seq data analysis. Raw reads were first subjected to adapter trimming using Cutadapt (RRID:SCR_011841) (Martin, 2011), followed by library complexity estimation using Preseq (RRID:SCR_018664), enabling projection of sequencing diversity and detection saturation. For peak detection, MACS2 (RRID:SCR_013291) (Zhang et al., 2008) was employed with merged input controls generated by aggregating input replicates corresponding to each IP group. To ensure consistency and reproducibility across biological replicates, replicate concordance was assessed using bamCompare from the deepTools suite (RRID:SCR_016366) (Ramirez et al., 2016). Further, deepTools was also utilized to generate normalized heatmaps depicting read pileups surrounding peak regions, providing insight into binding enrichment and distribution. Differential binding analysis was conducted using DESeq2 (RRID:SCR_015687) and ChIP-seq peaks with log2 Fold change >0.58 or <-0.58 and *P-*value ≤0.05 were considered significant for further analysis. For gene set Over Representation Analysis (ORA) for GO biological processes and significant KEGG pathways (RRID:SCR_018145), WEB-based GEne SeT Analysis Toolkit was used (RRID:SCR_006786) (Elizarraras et al., 2024). Raw and processed files have been deposited at the GEO database (see Supplementary file 7 for accession numbers).

### Reverse transcription, qPCR and ChIP-qPCR

RNA was isolated using the RNeasy Mini Kit (Qiagen), and reverse transcription was performed using the iScript cDNA Synthesis Kit (Bio-Rad, Cat#170-8891) according to the manufacturer’s instructions. qPCR and ChIP-qPCR were performed using iTaq Universal SYBR Green Supermix (Bio-Rad, Cat#1725121). For gene-expression qPCR, target gene expression was normalized to Gapdh expression. ChIP-qPCR enrichment was expressed as a percentage of input chromatin.

### Nuclear extracts and western blots

For nuclear protein extracts, nuclei were isolated using Buffer A [10 mM Hepes, 1.5 mM MgCl₂, 10 mM KCl], followed by nuclear disruption in Hepes lysis buffer [50 mM Hepes (pH 7.9), 150 mM NaCl, 0.5% Triton]. Both buffers were supplemented with EDTA-free protease inhibitors. The lysates were resuspended in 1× Laemmli buffer for subsequent immunoblotting. Lysates were loaded onto a 4-12% Bis-Tris gel and transferred to a PVDF membrane using the eBlot™ L1 fast wet transfer system (GenScript). The transferred proteins were incubated with blocking buffer (TBS [pH 8], 0.1% Tween 20, and 5% non-fat dry milk) for 1 hour at room temperature, followed by overnight immunoblotting with primary antibodies (see Supplementary file 7) at 4°C. Protein detection was performed using SuperSignal West Pico PLUS Chemiluminescent Substrates (Protein Biology, 34577).

### Confocal imaging and image processing

Whole mount colorimetric in situ embryos were imaged on a Zeiss Axioplan2 microscope. For whole mount FISH experiments, Ce3D++ was removed and confocal images were acquired on a Nikon A1R microscope using a 10x objective; NA = 0.4 and 3.3μm Z-sections. For HCR experiments in which embryos lacked an overt phenotype, embedding and image acquisition were performed without the knowledge of genotype. Tissue was subsequently recovered for genotyping. For midline sagittal sections of embryos processed for whole mount immunohistochemistry (see Figure 4B), the posterior third of the embryo was dissected, flat mounted and scanned from ventral to dorsal using a 25x silicon objective; NA=1.0 and 0.2μm Z-sections. All images were denoised and sharpened using NIS elements software (Nikon; RRID:SCR_014329) and image projections were analyzed using Fiji (RRID:SCR_002285) or Imaris software (v9.8-10.0.1, Oxford Instruments; RRID:SCR_007370). For optical cross sections through the primitive streak, the posterior third of the embryo was embedded vertically in 1% agarose on a glass bottom dish and confocal scanning was performed from posterior to anterior at 1μm Z-sections. A minimum of 3 embryos and 5 gastruloids were analyzed for each genotype per probe.

### Embryonic stem cell lines, culture and gastruloid assay

R1 ESCs (provided by Dr. Andras Nagy, Mount Sinai Hospital, Toronto) and A2LOX.Cre ESCs (provided by Dr. Michael Kyba, University of Minnesota, MN) were routinely cultured on mouse embryonic fibroblasts or feeder-free 0.1% gelatin coated dishes as described previously (Kennedy et al., 2016). *Wnt3a^ΔDE^* and *Sp5/8 dko* ESCs were generated from R1 ESCs using CRISpR-Cas9 targeting strategy (Figure 6-figure supplement 1). Serum and feeder-free ESCs were maintained in N2B27 media supplemented with LIF, CHIR99021, and PD0325901 (2i+LIF) on gelatin coated dishes. Before gastruloid experiments were initiated, ESCs were passaged at least 5 times in GMEM/10% FBS supplemented with non-essential amino acids, Glutamax, Sodium pyruvate, beta-mercaptoethanol and LIF as described previously (Beccari et al., 2018). ESCs were grown to 60-70% confluency and split using TrypLE select (Thermofisher Scientific, 12563029). Cells were washed twice with warm PBS (without CaCl2 and MgCl2) and resuspended in NDiff227 (Takara, Y40002) or ESGRO complete basal media (Millipore Sigma, SF002-500). Gastruloid formation was initiated with 300 cells/well in 96 well ultra-low attachment dishes (Corning, 3474). After 2 days, aggregates were treated with 150μl of warm media supplemented with 1μM CHIR (Tocris, 4423). Pre-warmed media (37^0^C) was exchanged on D3 and D4, and gastruloids were harvested on D5 using a cut 1000μl tip and fixed overnight in 4% PFA followed by dehydration in a methanol series the following morning. Gastruloids that failed to aggregate or elongate (15-20%) were excluded from experiments. All experiments with gastruloids were repeated at least 3 or more times.

### Luciferase assays

330ng of luciferase and 5ng of Renilla constructs were co-transfected into wildtype R1 (RRID:CVCL_2167) or A2Lox.Cre ES cells on Day 1 of culture. Cells were treated with CHIR+bFgf from Day 2 to Day 3. Luciferase assays were performed according to Dual Luciferase Assay Kit (Promega) instructions.

### Gene targeting

To generate knockout cell lines, the nucleofection of CRISpR-Cas9/gRNA ribonucleoprotein complexes and pMax-GFP constructs was prepared with the Amaxa^TM^ mouse ES cell nucleofector^TM^ Kit (Roche, VPH-1001) following the manufacturer’s instructions. The gRNA sequences are described in Supplementary file 7. The nucleofection of Cas9/gRNA complex and pMax-GFP cocktail solutions into 2X10^6^ cells was performed using the Amaxa Nucleofector (Lonza) in R1 ESCs (RRID:CVCL_2167). One day after nucleofection, GFP-positive cells were sorted using SH800 (Sony), and 3 X 10^5^ cells were plated onto a gelatin-coated 150mm dish. Colonies grown for 4-5days were picked and moved to a gelatin-coated 48 well plate. Genomic deletion was confirmed by Sanger sequencing (Psomagen, Rockville, MD).

### Data availability

Genomics data generated in this study have been deposited in the Gene Expression Omnibus (GEO) under accession numbers GSE252659 (*Sp5/8 dko* and *Wnt3a ko* RNA-seq), GSE291742 (Flag-Sp8 ChIP-seq), GSE293442 (Tbxt ChIP-seq), GSE296846 (D3 Flag-Sp5, Tcf7 and Tle ChIP-seq) and available to pubic. Accession numbers for all reused datasets are listed in Supplementary file 7. Source code has been provided for bulk-RNA seq analysis. Source data files have been provided for Figure 6-figure supplements 1A, 1C, 1D and Figure 7-figure supplement 1C.

### Use of AI tools in the manuscript

We used Claude to detect redundancies, missing references, reagents, and Research Resource Identifier, track changes and detect any grammatical errors. We reviewed, edited and verified all the content.

## Acknowledgements

We thank Erin Davies, NCI-Frederick for comments on the manuscript, Ms. Ruth Wolfe for excellent animal colony management, as well as the CCR sequencing facility. This work utilized the computational resources of the NIH HPC Biowulf cluster (https://hpc.nih.gov).

## References

Abu-Abed, S., Dolle, P., Metzger, D., Beckett, B., Chambon, P., Petkovich, M., 2001. The retinoic acid-metabolizing enzyme, CYP26A1, is essential for normal hindbrain patterning, vertebral identity, and development of posterior structures. Genes & development 15, 226–240.

Abu-Abed, S., Dolle, P., Metzger, D., Wood, C., MacLean, G., Chambon, P., Petkovich, M., 2003. Developing with lethal RA levels: genetic ablation of Rarg can restore the viability of mice lacking Cyp26a1. Development 130, 1449–1459.

Aires, R., de Lemos, L., Novoa, A., Jurberg, A.D., Mascrez, B., Duboule, D., Mallo, M., 2019. Tail Bud Progenitor Activity Relies on a Network Comprising Gdf11, Lin28, and Hox13 Genes. Developmental cell 48, 383–395 e388.

Aires, R., Dias, A., Mallo, M., 2018. Deconstructing the molecular mechanisms shaping the vertebrate body plan. Curr Opin Cell Biol 55, 81–86.

Amblard, I., Baranasic, D., Moyon, B., Lenhard, B., Metzis, V., 2024. A dual enhancer-silencer element ensures transient <em>Cdx2</em> expression during posterior body formation. bioRxiv, 2024.2004.2010.588878.

Amin, S., Neijts, R., Simmini, S., van Rooijen, C., Tan, S.C., Kester, L., van Oudenaarden, A., Creyghton, M.P., Deschamps, J., 2016. Cdx and T Brachyury Co-activate Growth Signaling in the Embryonic Axial Progenitor Niche. Cell reports 17, 3165–3177.

Anand, G.M., Megale, H.C., Murphy, S.H., Weis, T., Lin, Z., He, Y., Wang, X., Liu, J., Ramanathan, S., 2023. Controlling organoid symmetry breaking uncovers an excitable system underlying human axial elongation. Cell 186, 497–512 e423.

Anderson, M.J., Magidson, V., Kageyama, R., Lewandoski, M., 2020. Fgf4 maintains Hes7 levels critical for normal somite segmentation clock function. Elife 9.

Argelaguet, R., Lohoff, T., Li, J.G., Nakhuda, A., Drage, D., Krueger, F., Velten, L., Clark, S.J., Reik, W., 2022. Decoding gene regulation in the mouse embryo using single-cell multi-omics. bioRxiv, 2022.2006.2015.496239.

Arnold, S.J., Stappert, J., Bauer, A., Kispert, A., Herrmann, B.G., Kemler, R., 2000. Brachyury is a target gene of the Wnt/beta-catenin signaling pathway. Mechanisms of development 91, 249– 258.

Beccari, L., Moris, N., Girgin, M., Turner, D.A., Baillie-Johnson, P., Cossy, A.C., Lutolf, M.P., Duboule, D., Arias, A.M., 2018. Multi-axial self-organization properties of mouse embryonic stem cells into gastruloids. Nature 562, 272–276.

Beland, M., Pilon, N., Houle, M., Oh, K., Sylvestre, J.R., Prinos, P., Lohnes, D., 2004. Cdx1 autoregulation is governed by a novel Cdx1-LEF1 transcription complex. Molecular and cellular biology 24, 5028–5038.

Bell, S.M., Schreiner, C.M., Waclaw, R.R., Campbell, K., Potter, S.S., Scott, W.J., 2003. Sp8 is crucial for limb outgrowth and neuropore closure. Proceedings of the National Academy of Sciences of the United States of America 100, 12195–12200.

Binagui-Casas, A., Dias, A., Guillot, C., Metzis, V., Saunders, D., 2021. Building consensus in neuromesodermal research: Current advances and future biomedical perspectives. Curr Opin Cell Biol 73, 133–140.

Blassberg, R., Patel, H., Watson, T., Gouti, M., Metzis, V., Delas, M.J., Briscoe, J., 2022. Sox2 levels regulate the chromatin occupancy of WNT mediators in epiblast progenitors responsible for vertebrate body formation. Nature cell biology 24, 633–644.

Bou-Rouphael, J., Durand, B.C., 2021. T-Cell Factors as Transcriptional Inhibitors: Activities and Regulations in Vertebrate Head Development. Front Cell Dev Biol 9, 784998.

Boulet, A.M., Capecchi, M.R., 2012. Signaling by FGF4 and FGF8 is required for axial elongation of the mouse embryo. Developmental biology 371, 235–245.

Boylan, M., Anderson, M.J., Ornitz, D.M., Lewandoski, M., 2020. The Fgf8 subfamily (Fgf8, Fgf17 and Fgf18) is required for closure of the embryonic ventral body wall. Development 147.

Cambray, N., Wilson, V., 2002. Axial progenitors with extensive potency are localised to the mouse chordoneural hinge. Development 129, 4855–4866.

Cambray, N., Wilson, V., 2007. Two distinct sources for a population of maturing axial progenitors. Development 134, 2829–2840.

Chalamalasetty, R.B., Dunty, W.C., Jr., Biris, K.K., Ajima, R., Iacovino, M., Beisaw, A., Feigenbaum, L., Chapman, D.L., Yoon, J.K., Kyba, M., Yamaguchi, T.P., 2011. The Wnt3a/beta-catenin target gene Mesogenin1 controls the segmentation clock by activating a Notch signalling program. Nature communications 2, 390.

Chalamalasetty, R.B., Garriock, R.J., Dunty, W.C., Jr., Kennedy, M.W., Jailwala, P., Si, H., Yamaguchi, T.P., 2014. Mesogenin 1 is a master regulator of paraxial presomitic mesoderm differentiation. Development 141, 4285–4297.

Chawengsaksophak, K., de Graaff, W., Rossant, J., Deschamps, J., Beck, F., 2004. Cdx2 is essential for axial elongation in mouse development. Proceedings of the National Academy of Sciences of the United States of America 101, 7641–7645.

Chawengsaksophak, K., James, R., Hammond, V.E., Kontgen, F., Beck, F., 1997. Homeosis and intestinal tumours in Cdx2 mutant mice. Nature 386, 84–87.

Chodaparambil, J.V., Pate, K.T., Hepler, M.R., Tsai, B.P., Muthurajan, U.M., Luger, K., Waterman, M.L., Weis, W.I., 2014. Molecular functions of the TLE tetramerization domain in Wnt target gene repression. The EMBO journal 33, 719–731.

Choi, H.M.T., Schwarzkopf, M., Fornace, M.E., Acharya, A., Artavanis, G., Stegmaier, J., Cunha, A., Pierce, N.A., 2018. Third-generation in situ hybridization chain reaction: multiplexed, quantitative, sensitive, versatile, robust. Development 145.

Clevers, H., Loh, K.M., Nusse, R., 2014. Stem cell signaling. An integral program for tissue renewal and regeneration: Wnt signaling and stem cell control. Science 346, 1248012.

Crossley, P.H., Martin, G.R., 1995. The mouse Fgf8 gene encodes a family of polypeptides and is expressed in regions that direct outgrowth and patterning in the developing embryo. Development 121, 439–451.

Cunningham, T.J., Kumar, S., Yamaguchi, T.P., Duester, G., 2015. Wnt8a and Wnt3a cooperate in the axial stem cell niche to promote mammalian body axis extension. Developmental dynamics : an official publication of the American Association of Anatomists 244, 797–807.

Dailey, S.C., Kozmikova, I., Somorjai, I.M.L., 2017. Amphioxus Sp5 is a member of a conserved Specificity Protein complement and is modulated by Wnt/beta-catenin signalling. Int J Dev Biol 61, 723–732.

Deschamps, J., Duboule, D., 2017. Embryonic timing, axial stem cells, chromatin dynamics, and the Hox clock. Genes & development 31, 1406–1416.

Dobin, A., Davis, C.A., Schlesinger, F., Drenkow, J., Zaleski, C., Jha, S., Batut, P., Chaisson, M., Gingeras, T.R., 2013. STAR: ultrafast universal RNA-seq aligner. Bioinformatics 29, 15–21.

Dunty, W.C., Jr., Biris, K.K., Chalamalasetty, R.B., Taketo, M.M., Lewandoski, M., Yamaguchi, T.P., 2008. Wnt3a/beta-catenin signaling controls posterior body development by coordinating mesoderm formation and segmentation. Development 135, 85–94.

Dunty, W.C., Jr., Kennedy, M.W., Chalamalasetty, R.B., Campbell, K., Yamaguchi, T.P., 2014. Transcriptional profiling of Wnt3a mutants identifies Sp transcription factors as essential effectors of the Wnt/beta-catenin pathway in neuromesodermal stem cells. PloS one 9, e87018.

Elizarraras, J.M., Liao, Y., Shi, Z., Zhu, Q., Pico, A.R., Zhang, B., 2024. WebGestalt 2024: faster gene set analysis and new support for metabolomics and multi-omics. Nucleic acids research 52, W415–W421.

Frazer, K.A., Pachter, L., Poliakov, A., Rubin, E.M., Dubchak, I., 2004. VISTA: computational tools for comparative genomics. Nucleic acids research 32, W273–279.

Galceran, J., Farinas, I., Depew, M.J., Clevers, H., Grosschedl, R., 1999. Wnt3a-/--like phenotype and limb deficiency in Lef1(-/-)Tcf1(-/-) mice. Genes & development 13, 709–717.

Galceran, J., Hsu, S.C., Grosschedl, R., 2001. Rescue of a Wnt mutation by an activated form of LEF-1: regulation of maintenance but not initiation of Brachyury expression. Proceedings of the National Academy of Sciences of the United States of America 98, 8668–8673.

Garriock, R.J., Chalamalasetty, R.B., Kennedy, M.W., Canizales, L.C., Lewandoski, M., Yamaguchi, T.P., 2015. Lineage tracing of neuromesodermal progenitors reveals novel Wnt-dependent roles in trunk progenitor cell maintenance and differentiation. Development 142, 1628–1638.

Garriock, R.J., Chalamalasetty, R.B., Zhu, J., Kennedy, M.W., Kumar, A., Mackem, S., Yamaguchi, T.P., 2020. A dorsal-ventral gradient of Wnt3a/beta-catenin signals control hindgut extension and colon formation. Development.

Gautam, S., Fenner, J.L., Wang, B., Range, R.C., 2024. Evolutionarily conserved Wnt/Sp5 signaling is critical for anterior-posterior axis patterning in sea urchin embryos. iScience 27, 108616.

Gearing, L.J., Cumming, H.E., Chapman, R., Finkel, A.M., Woodhouse, I.B., Luu, K., Gould, J.A., Forster, S.C., Hertzog, P.J., 2019. CiiiDER: A tool for predicting and analysing transcription factor binding sites. PloS one 14, e0215495.

Gouti, M., Delile, J., Stamataki, D., Wymeersch, F.J., Huang, Y., Kleinjung, J., Wilson, V., Briscoe, J., 2017. A Gene Regulatory Network Balances Neural and Mesoderm Specification during Vertebrate Trunk Development. Developmental cell 41, 243–261 e247.

Gouti, M., Tsakiridis, A., Wymeersch, F.J., Huang, Y., Kleinjung, J., Wilson, V., Briscoe, J., 2014. In vitro generation of neuromesodermal progenitors reveals distinct roles for wnt signalling in the specification of spinal cord and paraxial mesoderm identity. PLoS biology 12, e1001937.

Greco, T.L., Takada, S., Newhouse, M.M., McMahon, J.A., McMahon, A.P., Camper, S.A., 1996. Analysis of the vestigial tail mutation demonstrates that Wnt-3a gene dosage regulates mouse axial development. Genes & development 10, 313–324.

Guibentif, C., Griffiths, J.A., Imaz-Rosshandler, I., Ghazanfar, S., Nichols, J., Wilson, V., Gottgens, B., Marioni, J.C., 2021. Diverse Routes toward Early Somites in the Mouse Embryo. Developmental cell 56, 141–153 e146.

Guo, Q., Kim, A., Li, B., Ransick, A., Bugacov, H., Chen, X., Lindstrom, N., Brown, A., Oxburgh, L., Ren, B., McMahon, A.P., 2021. A beta-catenin-driven switch in TCF/LEF transcription factor binding to DNA target sites promotes commitment of mammalian nephron progenitor cells. Elife 10.

Harrison, S.M., Houzelstein, D., Dunwoodie, S.L., Beddington, R.S., 2000. Sp5, a new member of the Sp1 family, is dynamically expressed during development and genetically interacts with Brachyury. Developmental biology 227, 358–372.

Henrique, D., Abranches, E., Verrier, L., Storey, K.G., 2015. Neuromesodermal progenitors and the making of the spinal cord. Development 142, 2864–2875.

Hoffman, J.A., Wu, C.I., Merrill, B.J., 2013. Tcf7l1 prepares epiblast cells in the gastrulating mouse embryo for lineage specification. Development 140, 1665–1675.

Hogan, B., Hogan, B., 1994. Manipulating the mouse embryo : a laboratory manual, 2nd ed. Cold Spring Harbor Laboratory Press, Plainview, N.Y.

Hovanes, K., Li, T.W., Munguia, J.E., Truong, T., Milovanovic, T., Lawrence Marsh, J., Holcombe, R.F., Waterman, M.L., 2001. Beta-catenin-sensitive isoforms of lymphoid enhancer factor-1 are selectively expressed in colon cancer. Nature genetics 28, 53–57.

Huggins, I.J., Bos, T., Gaylord, O., Jessen, C., Lonquich, B., Puranen, A., Richter, J., Rossdam, C., Brafman, D., Gaasterland, T., Willert, K., 2017. The WNT target SP5 negatively regulates WNT transcriptional programs in human pluripotent stem cells. Nature communications 8, 1034.

Iacovino, M., Bosnakovski, D., Fey, H., Rux, D., Bajwa, G., Mahen, E., Mitanoska, A., Xu, Z., Kyba, M., 2011. Inducible cassette exchange: a rapid and efficient system enabling conditional gene expression in embryonic stem and primary cells. Stem cells 29, 1580–1588.

Jurberg, A.D., Aires, R., Varela-Lasheras, I., Novoa, A., Mallo, M., 2013. Switching axial progenitors from producing trunk to tail tissues in vertebrate embryos. Developmental cell 25, 451–462.

Kawakami, Y., Esteban, C.R., Matsui, T., Rodriguez-Leon, J., Kato, S., Belmonte, J.C., 2004. Sp8 and Sp9, two closely related buttonhead-like transcription factors, regulate Fgf8 expression and limb outgrowth in vertebrate embryos. Development 131, 4763–4774.

Kennedy, M.W., Chalamalasetty, R.B., Thomas, S., Garriock, R.J., Jailwala, P., Yamaguchi, T.P., 2016. Sp5 and Sp8 recruit beta-catenin and Tcf1-Lef1 to select enhancers to activate Wnt target gene transcription. Proceedings of the National Academy of Sciences of the United States of America 113, 3545–3550.

Koch, F., Scholze, M., Wittler, L., Schifferl, D., Sudheer, S., Grote, P., Timmermann, B., Macura, K., Herrmann, B.G., 2017. Antagonistic Activities of Sox2 and Brachyury Control the Fate Choice of Neuro-Mesodermal Progenitors. Developmental cell 42, 514–526 e517.

Liang, Y., Koche, R., Chalamalasetty, R.B., Stephen, D.N., Kennedy, M.W., Lao, Z., Pang, Y., Kuo, Y.Y., Lee, M., Lobo, F.P., Huang, X., Hadjantonakis, A.K., Yamaguchi, T.P., Anderson, K.V., Joyner, A.L., 2025. Transcription factors SP5 and SP8 drive primary cilia formation in mammalian embryos. Science 389, eadt5663.

Love, M.I., Huber, W., Anders, S., 2014. Moderated estimation of fold change and dispersion for RNA-seq data with DESeq2. Genome biology 15, 550.

MacMurray, A., Shin, H.S., 1988. The antimorphic nature of the Tc allele at the mouse T locus. Genetics 120, 545–550.

Martin, B.L., Kimelman, D., 2010. Brachyury establishes the embryonic mesodermal progenitor niche. Genes & development 24, 2778–2783.

Martin, B.L., Kimelman, D., 2012. Canonical Wnt signaling dynamically controls multiple stem cell fate decisions during vertebrate body formation. Developmental cell 22, 223–232.

Martin, M., 2011. Cutadapt removes adapter sequences from high-throughput sequencing reads. 2011 17, 3.

Maruoka, Y., Ohbayashi, N., Hoshikawa, M., Itoh, N., Hogan, B.L., Furuta, Y., 1998. Comparison of the expression of three highly related genes, Fgf8, Fgf17 and Fgf18, in the mouse embryo. Mechanisms of development 74, 175–177.

Mohanty, S., Lekven, A.C., 2025. Divergent functions of the evolutionarily conserved, yet seemingly dispensable, Wnt target, sp5. Differentiation; research in biological diversity 141, 100829.

Naiche, L.A., Holder, N., Lewandoski, M., 2011. FGF4 and FGF8 comprise the wavefront activity that controls somitogenesis. Proceedings of the National Academy of Sciences of the United States of America 108, 4018–4023.

Nowotschin, S., Ferrer-Vaquer, A., Concepcion, D., Papaioannou, V.E., Hadjantonakis, A.K., 2012. Interaction of Wnt3a, Msgn1 and Tbx6 in neural versus paraxial mesoderm lineage commitment and paraxial mesoderm differentiation in the mouse embryo. Developmental biology 367, 1–14.

Perantoni, A.O., Timofeeva, O., Naillat, F., Richman, C., Pajni-Underwood, S., Wilson, C., Vainio, S., Dove, L.F., Lewandoski, M., 2005. Inactivation of FGF8 in early mesoderm reveals an essential role in kidney development. Development 132, 3859–3871.

Pijuan-Sala, B., Griffiths, J.A., Guibentif, C., Hiscock, T.W., Jawaid, W., Calero-Nieto, F.J., Mulas, C., Ibarra-Soria, X., Tyser, R.C.V., Ho, D.L.L., Reik, W., Srinivas, S., Simons, B.D., Nichols, J., Marioni, J.C., Gottgens, B., 2019. A single-cell molecular map of mouse gastrulation and early organogenesis. Nature 566, 490–495.

Ramakrishnan, A.B., Cadigan, K.M., 2017. Wnt target genes and where to find them. F1000Res 6, 746.

Ramakrishnan, A.B., Chen, L., Burby, P.E., Cadigan, K.M., 2021. Wnt target enhancer regulation by a CDX/TCF transcription factor collective and a novel DNA motif. Nucleic acids research 49, 8625–8641.

Ramakrishnan, A.B., Sinha, A., Fan, V.B., Cadigan, K.M., 2018. The Wnt Transcriptional Switch: TLE Removal or Inactivation? BioEssays : news and reviews in molecular, cellular and developmental biology 40.

Ramirez, F., Ryan, D.P., Gruning, B., Bhardwaj, V., Kilpert, F., Richter, A.S., Heyne, S., Dundar, F., Manke, T., 2016. deepTools2: a next generation web server for deep-sequencing data analysis. Nucleic acids research 44, W160–165.

Rodrigo Albors, A., Halley, P.A., Storey, K.G., 2018. Lineage tracing of axial progenitors using Nkx1-2CreER(T2) mice defines their trunk and tail contributions. Development 145.

Row, R.H., Pegg, A., Kinney, B.A., Farr, G.H., 3rd, Maves, L., Lowell, S., Wilson, V., Martin, B.L., 2018. BMP and FGF signaling interact to pattern mesoderm by controlling basic helix-loop-helix transcription factor activity. Elife 7.

Sakai, Y., Meno, C., Fujii, H., Nishino, J., Shiratori, H., Saijoh, Y., Rossant, J., Hamada, H., 2001. The retinoic acid-inactivating enzyme CYP26 is essential for establishing an uneven distribution of retinoic acid along the anterio-posterior axis within the mouse embryo. Genes & development 15, 213–225.

Santos, A.J.M., Lo, Y.H., Mah, A.T., Kuo, C.J., 2018. The Intestinal Stem Cell Niche: Homeostasis and Adaptations. Trends Cell Biol 28, 1062–1078.

Savory, J.G., Bouchard, N., Pierre, V., Rijli, F.M., De Repentigny, Y., Kothary, R., Lohnes, D., 2009. Cdx2 regulation of posterior development through non-Hox targets. Development 136, 4099–4110.

Sayols, S., 2023. rrvgo: a Bioconductor package for interpreting lists of Gene Ontology terms. MicroPubl Biol 2023.

Schnirman, R.E., Kuo, S.J., Kelly, R.C., Yamaguchi, T.P., 2023. The role of Wnt signaling in the development of the epiblast and axial progenitors. Curr Top Dev Biol 153, 145–180.

Shy, B.R., Wu, C.I., Khramtsova, G.F., Zhang, J.Y., Olopade, O.I., Goss, K.H., Merrill, B.J., 2013. Regulation of Tcf7l1 DNA binding and protein stability as principal mechanisms of Wnt/beta-catenin signaling. Cell reports 4, 1–9.

Soneson, C., Love, M.I., Robinson, M.D., 2015. Differential analyses for RNA-seq: transcript-level estimates improve gene-level inferences. F1000Res 4, 1521.

Steventon, B., Martinez Arias, A., 2017. Evo-engineering and the cellular and molecular origins of the vertebrate spinal cord. Developmental biology 432, 3–13.

Stott, D., Kispert, A., Herrmann, B.G., 1993. Rescue of the tail defect of Brachyury mice. Genes & development 7, 197–203.

Sudheer, S., Liu, J., Marks, M., Koch, F., Anurin, A., Scholze, M., Senft, A.D., Wittler, L., Macura, K., Grote, P., Herrmann, B.G., 2016. Different Concentrations of FGF Ligands, FGF2 or FGF8 Determine Distinct States of WNT-Induced Presomitic Mesoderm. Stem cells 34, 1790–1800.

Takada, S., Stark, K.L., Shea, M.J., Vassileva, G., McMahon, J.A., McMahon, A.P., 1994. Wnt-3a regulates somite and tailbud formation in the mouse embryo. Genes & development 8, 174– 189.

Tewari, A.G., Owen, J.H., Petersen, C.P., Wagner, D.E., Reddien, P.W., 2019. A small set of conserved genes, including sp5 and Hox, are activated by Wnt signaling in the posterior of planarians and acoels. PLoS genetics 15, e1008401.

Tran, T.H.N., Takada, R., Krayukhina, E., Maruno, T., Mii, Y., Uchiyama, S., Takada, S., 2024. Soluble Frizzled-related proteins promote exosome-mediated Wnt re-secretion. Commun Biol 7, 254.

Tsakiridis, A., Huang, Y., Blin, G., Skylaki, S., Wymeersch, F., Osorno, R., Economou, C., Karagianni, E., Zhao, S., Lowell, S., Wilson, V., 2014. Distinct Wnt-driven primitive streak-like populations reflect in vivo lineage precursors. Development 141, 1209–1221.

Tzouanacou, E., Wegener, A., Wymeersch, F.J., Wilson, V., Nicolas, J.F., 2009. Redefining the progression of lineage segregations during mammalian embryogenesis by clonal analysis. Developmental cell 17, 365–376.

van de Ven, C., Bialecka, M., Neijts, R., Young, T., Rowland, J.E., Stringer, E.J., Van Rooijen, C., Meijlink, F., Novoa, A., Freund, J.N., Mallo, M., Beck, F., Deschamps, J., 2011. Concerted involvement of Cdx/Hox genes and Wnt signaling in morphogenesis of the caudal neural tube and cloacal derivatives from the posterior growth zone. Development 138, 3451–3462.

van den Akker, E., Forlani, S., Chawengsaksophak, K., de Graaff, W., Beck, F., Meyer, B.I., Deschamps, J., 2002. Cdx1 and Cdx2 have overlapping functions in anteroposterior patterning and posterior axis elongation. Development 129, 2181–2193.

van den Brink, S.C., Alemany, A., van Batenburg, V., Moris, N., Blotenburg, M., Vivie, J., Baillie-Johnson, P., Nichols, J., Sonnen, K.F., Martinez Arias, A., van Oudenaarden, A., 2020. Single-cell and spatial transcriptomics reveal somitogenesis in gastruloids. Nature 582, 405–409.

van Nes, J., de Graaff, W., Lebrin, F., Gerhard, M., Beck, F., Deschamps, J., 2006. The Cdx4 mutation affects axial development and reveals an essential role of Cdx genes in the ontogenesis of the placental labyrinth in mice. Development 133, 419–428.

van Tienen, L.M., Mieszczanek, J., Fiedler, M., Rutherford, T.J., Bienz, M., 2017. Constitutive scaffolding of multiple Wnt enhanceosome components by Legless/BCL9. Elife 6.

Veenvliet, J.V., Bolondi, A., Kretzmer, H., Haut, L., Scholze-Wittler, M., Schifferl, D., Koch, F., Guignard, L., Kumar, A.S., Pustet, M., Heimann, S., Buschow, R., Wittler, L., Timmermann, B., Meissner, A., Herrmann, B.G., 2020. Mouse embryonic stem cells self-organize into trunk-like structures with neural tube and somites. Science 370.

Vogg, M.C., Beccari, L., Iglesias Olle, L., Rampon, C., Vriz, S., Perruchoud, C., Wenger, Y., Galliot, B., 2019. An evolutionarily-conserved Wnt3/beta-catenin/Sp5 feedback loop restricts head organizer activity in Hydra. Nature communications 10, 312.

Wahl, M.B., Deng, C., Lewandoski, M., Pourquie, O., 2007. FGF signaling acts upstream of the NOTCH and WNT signaling pathways to control segmentation clock oscillations in mouse somitogenesis. Development 134, 4033–4041.

Wang, L., Wang, S., Li, W., 2012. RSeQC: quality control of RNA-seq experiments. Bioinformatics 28, 2184–2185.

Weidinger, G., Thorpe, C.J., Wuennenberg-Stapleton, K., Ngai, J., Moon, R.T., 2005. The Sp1-related transcription factors sp5 and sp5-like act downstream of Wnt/beta-catenin signaling in mesoderm and neuroectoderm patterning. Current biology : CB 15, 489–500.

Wilson, V., Olivera-Martinez, I., Storey, K.G., 2009. Stem cells, signals and vertebrate body axis extension. Development 136, 1591–1604.

Wingett, S.W., Andrews, S., 2018. FastQ Screen: A tool for multi-genome mapping and quality control. F1000Res 7, 1338.

Wood, D.E., Lu, J., Langmead, B., 2019. Improved metagenomic analysis with Kraken 2. Genome biology 20, 257.

Wu, C.I., Hoffman, J.A., Shy, B.R., Ford, E.M., Fuchs, E., Nguyen, H., Merrill, B.J., 2012. Function of Wnt/beta-catenin in counteracting Tcf3 repression through the Tcf3-beta-catenin interaction. Development 139, 2118–2129.

Wymeersch, F.J., Huang, Y., Blin, G., Cambray, N., Wilkie, R., Wong, F.C., Wilson, V., 2016. Position-dependent plasticity of distinct progenitor types in the primitive streak. Elife 5, e10042.

Wymeersch, F.J., Skylaki, S., Huang, Y., Watson, J.A., Economou, C., Marek-Johnston, C., Tomlinson, S.R., Wilson, V., 2019. Transcriptionally dynamic progenitor populations organised around a stable niche drive axial patterning. Development 146.

Xiang, Y., Zhang, Y., Xu, Q., Zhou, C., Liu, B., Du, Z., Zhang, K., Zhang, B., Wang, X., Gayen, S., Liu, L., Wang, Y., Li, Y., Wang, Q., Kalantry, S., Li, L., Xie, W., 2020. Epigenomic analysis of gastrulation identifies a unique chromatin state for primed pluripotency. Nature genetics 52, 95– 105.

Yamaguchi, T.P., Rossant, J., 1995. Fibroblast growth factors in mammalian development. Curr Opin Genet Dev 5, 485–491.

Yamaguchi, T.P., Takada, S., Yoshikawa, Y., Wu, N., McMahon, A.P., 1999. T (Brachyury) is a direct target of Wnt3a during paraxial mesoderm specification. Genes & development 13, 3185– 3190.

Yang, X., Hu, B., Liao, J., Qiao, Y., Chen, Y., Qian, Y., Feng, S., Yu, F., Dong, J., Hou, Y., Xu, H., Wang, R., Peng, G., Li, J., Tang, F., Jing, N., 2019. Distinct enhancer signatures in the mouse gastrula delineate progressive cell fate continuum during embryo development. Cell research 29, 911–926.

Yeo, G.H.T., Lin, L., Qi, C.Y., Cha, M., Gifford, D.K., Sherwood, R.I., 2020. A Multiplexed Barcodelet Single-Cell RNA-Seq Approach Elucidates Combinatorial Signaling Pathways that Drive ESC Differentiation. Cell stem cell 26, 938–950 e936.

Yi, F., Pereira, L., Hoffman, J.A., Shy, B.R., Yuen, C.M., Liu, D.R., Merrill, B.J., 2011. Opposing effects of Tcf3 and Tcf1 control Wnt stimulation of embryonic stem cell self-renewal. Nature cell biology 13, 762–770.

Yoshikawa, Y., Fujimori, T., McMahon, A.P., Takada, S., 1997. Evidence that absence of Wnt-3a signaling promotes neuralization instead of paraxial mesoderm development in the mouse. Developmental biology 183, 234–242.

Young, T., Rowland, J.E., van de Ven, C., Bialecka, M., Novoa, A., Carapuco, M., van Nes, J., de Graaff, W., Duluc, I., Freund, J.N., Beck, F., Mallo, M., Deschamps, J., 2009. Cdx and Hox genes differentially regulate posterior axial growth in mammalian embryos. Developmental cell 17, 516–526.

Yu, G., Wang, L.G., Han, Y., He, Q.Y., 2012. clusterProfiler: an R package for comparing biological themes among gene clusters. OMICS 16, 284–287.

Zembrzycki, A., Griesel, G., Stoykova, A., Mansouri, A., 2007. Genetic interplay between the transcription factors Sp8 and Emx2 in the patterning of the forebrain. Neural Dev 2, 8.

Zhang, Y., Liu, T., Meyer, C.A., Eeckhoute, J., Johnson, D.S., Bernstein, B.E., Nusbaum, C., Myers, R.M., Brown, M., Li, W., Liu, X.S., 2008. Model-based analysis of ChIP-Seq (MACS). Genome biology 9, R137.

Zhu, Y., Lohnes, D., 2022. Regulation of axial elongation by Cdx. Developmental biology 483, 118–127.

